# CXCR6⁺ CD127⁻ Tr1 Cells Balance Immunity and Persistence in Plasmodium falciparum Infection

**DOI:** 10.1101/2025.09.16.676649

**Authors:** Jason Nideffer, Florian Bach, Steven Strubbe, Luis Lopez, Maato Zedi, Felistas Nankya, Jessica Briggs, Kattria van der Ploeg, Kenneth Musinguzi, Soyeon Kim, Aracely Garcia Romero, Arefin Keya, Kylie Camanag, Savannah Lewis, Muhammad Abdelbasset, Bing Wang, Allison Boss, Evelyn Nansubuga, Joaniter I Nankabirwa, Emmanuel Arinaitwe, Saki Takahashi, Grant Dorsey, Bryan Greenhouse, Isabel Rodriguez-Barraquer, Moses R Kamya, Rosa Bacchetta, Isaac Ssewanyana, Ashraful Haque, Maria Grazia Roncarolo, Prasanna Jagannathan

## Abstract

*Plasmodium falciparum* (*Pf*) induces the clonal expansion of antigen-specific type 1 regulatory T (Tr1) cells with a capacity for long-term memory. Tr1 cells comprise nearly 90% of the *Pf* blood stage antigen-specific CD4+ T cell pool in children. Though, whether Tr1 cells contribute to protection from malaria remains undetermined. To address this critical gap in knowledge, we first performed scRNAseq on gated cell populations to validate CXCR6+ CD127- as new phenotypic markers to enrich for bona-fide Tr1 cells. Importantly, these Tr1 cells potently suppressed the proliferation of other CD4+ T cells *in vitro* via IL-10 secretion. Among children living in malaria-endemic Uganda, CXCR6+ CD127- Tr1 cells were the dominant responding subset to *Pf*-infected red blood cell stimulation *in vitro*. They also rapidly expanded following malaria and expressed IL-10 and IFNg during infection *in vivo*. Tr1 abundance correlated with plasma concentrations of granzyme A, IFNg, IL-10, and LAG3, suggesting that these cells act systemically. Higher CXCR6+ CD127- Tr1 cell frequencies correlated with a lower probability of symptoms given parasitemia but were also associated with delayed parasite clearance among untreated, asymptomatic children. These data suggest that Tr1 cells help mediate clinical immunity to malaria but may also facilitate parasite persistence through mechanisms of immune regulation.

## Introduction

Malaria is a life-threatening disease, particularly among children under five years of age living in malaria-endemic settings^1^. Although these children typically never develop protection from infection^2^, they eventually, through repeated exposure, develop “clinical immunity”—the ability to tolerate parasitemia without becoming sick^3^. It is generally thought that regulatory immune responses are critically important for attenuating harmful inflammation to support health despite parasitemia among clinically immune individuals^4^. While these regulatory responses may be beneficial in the setting of clinical immunity, it is possible that learned regulatory responses resulting from repeated natural infection may hinder efforts to clear infection, and may also explain reduced vaccine efficacy in malaria endemic, high-transmission regions^5^.

We recently demonstrated that the *Pf*-specific CD4+ T cell response is dominated by Type 1 regulatory T (Tr1) cells^6^, a subset of memory CD4+ T cells that lack FOXP3 but regulate immune responses through expression of IL-10 and coinhibitory receptors^7^. These Tr1 cells responded transcriptionally to *in vitro* stimulation with *Plasmodium* antigens; they clonally expanded following malaria; and, they demonstrated long-term memory with clonal fidelity—a strong, consistent bias among clones for the Tr1 identity over time^6^. Given the regulatory functions that have been attributed to Tr1 cells in many contexts^8^, including a recently described capacity to inhibit tumor immunity^9^, Tr1 cells are hypothesized to play a causal role in dampened inflammatory responses intrinsic to antimalarial clinical immunity^4^. However, the function and clinical relevance of Tr1 cells in the context of human malaria has not been determined.

To better understand the role of Tr1 cells in the acquisition of clinical and vaccine induced immunity, cell surface markers that enrich for malaria-specific Tr1 cells are essential. Prior studies have suggested the use of certain markers to identify Tr1 cells in peripheral blood of healthy individuals^10^. The cell surface markers CD49b and lymphocyte activation gene 3 (LAG-3) were transcriptionally enriched among IL-10 producing expanded human T cell clones. This gating strategy was then shown to enrich for CD4+ T cells that demonstrated suppressive functions *in vitro* and in mouse models of intestinal inflammation and helminth infection^10^. More recently, CCR5+ PD-1+ was shown to enrich for CD4+ T cells that suppress colitis in mice and that co-express IL-10 and IFNγ in the human gut^11^. While both of these surface marker-defined populations enrich for suppressive CD4+ T cells in different contexts, it is unclear whether these definitions can be used to identify malaria-specific Tr1 cells.

Here, we utilized single cell RNA sequencing to identify putative cell surface markers for Tr1 cells in the blood of children living in malaria-endemic Uganda and establish that CXCR6+ CD127-memory CD4+ T cells enrich for high-confidence, bona fide Tr1 cells in this setting. We further demonstrate that CXCR6+ CD127-Tr1 cells are functionally suppressive, respond to *Plasmodium* antigens *in vitro* and *in vivo*, and correlate with systemic responses to malaria. Lastly, we demonstrate that CXCR6+ CD127-Tr1 cells correlate with clinical outcomes, including a lower probability of symptoms given infection, as well as a prolonged time to infection clearance among children with asymptomatic, untreated infections, consistent with a role for these cells in mediating clinical immunity to malaria through their regulation of pathological immune responses.

## Results

### Validation of CXCR6+ CD127-as a gating strategy to enrich for malaria-specific Tr1 cells

We previously used single-cell RNA sequencing (scRNAseq) to identify Tr1 cells in the periphery of malaria-exposed Ugandan children (Fig. 1A). These cells expressed key Tr1-associated genes (*IL10*, *LAG3*, *CTLA4*, *PDCD1*, *MAF*, *PRDM1*, *GZMA*, *GZMK*); they were distinct from *FOXP3*-expressing Tregs; and, they demonstrated long-term memory with clonal fidelity (maintaining their Tr1 identity) upon recall^6^. These Tr1 cells expressed *CXCR6* and had low to no expression of *IL7R*—a pattern that appeared relatively unique among memory CD4+ T cells (Fig. 1B). We thus performed flow cytometric analysis for these markers, gating putative Tr1 cells as CXCR6+ CD127-(Fig. 1C), and quantified their frequencies across samples that were also analyzed by scRNAseq^6^. CXCR6+ CD127-frequencies measured by flow cytometry strongly correlated with Tr1 frequencies determined by scRNAseq (Fig. 1D). To validate this gating strategy, we then sorted memory CD4+ T cells from the blood of four Ugandan children based on their expression of CXCR6 and CD127 and performed scRNAseq (Fig. 1E). Merging these data with scRNAseq data from our prior study, we found that CXCR6+ CD127-cells were highly enriched for Tr1 cells (Fig. 1F). By quantifying the relative abundance of memory CD4+ T cells that are Tr1 or non-Tr1 and that fall inside and outside of the CXCR6+ CD127-gate (Fig. S1), we determined this gating strategy to be ∼80% sensitive and ∼99% specific for the identification of Tr1 cells among these individuals (Fig. 1G). The largest contaminating population among CXCR6+ CD127-memory CD4+ T cells (<10%) was FOXP3-expressing Tregs (Fig. S2A), consistent with conventional Tregs also typically lacking or lowly expressing CD127^12^. When considering the transcriptional heterogeneity that exists within the Tr1 subset^6^, this gating strategy captured 98.8% of effector Tr1 cells, 84.5% of memory Tr1 cells, and 99.3% of naïve-like Tr1 cells (Fig. S2B-D). However, activated Tr1 cells (and some memory Tr1 cells) downregulate CXCR6 (Fig. S2E), suggesting that CXCR6+ CD127-may not capture all Tr1 cell subsets with the same efficacy.

**Fig. 1.**
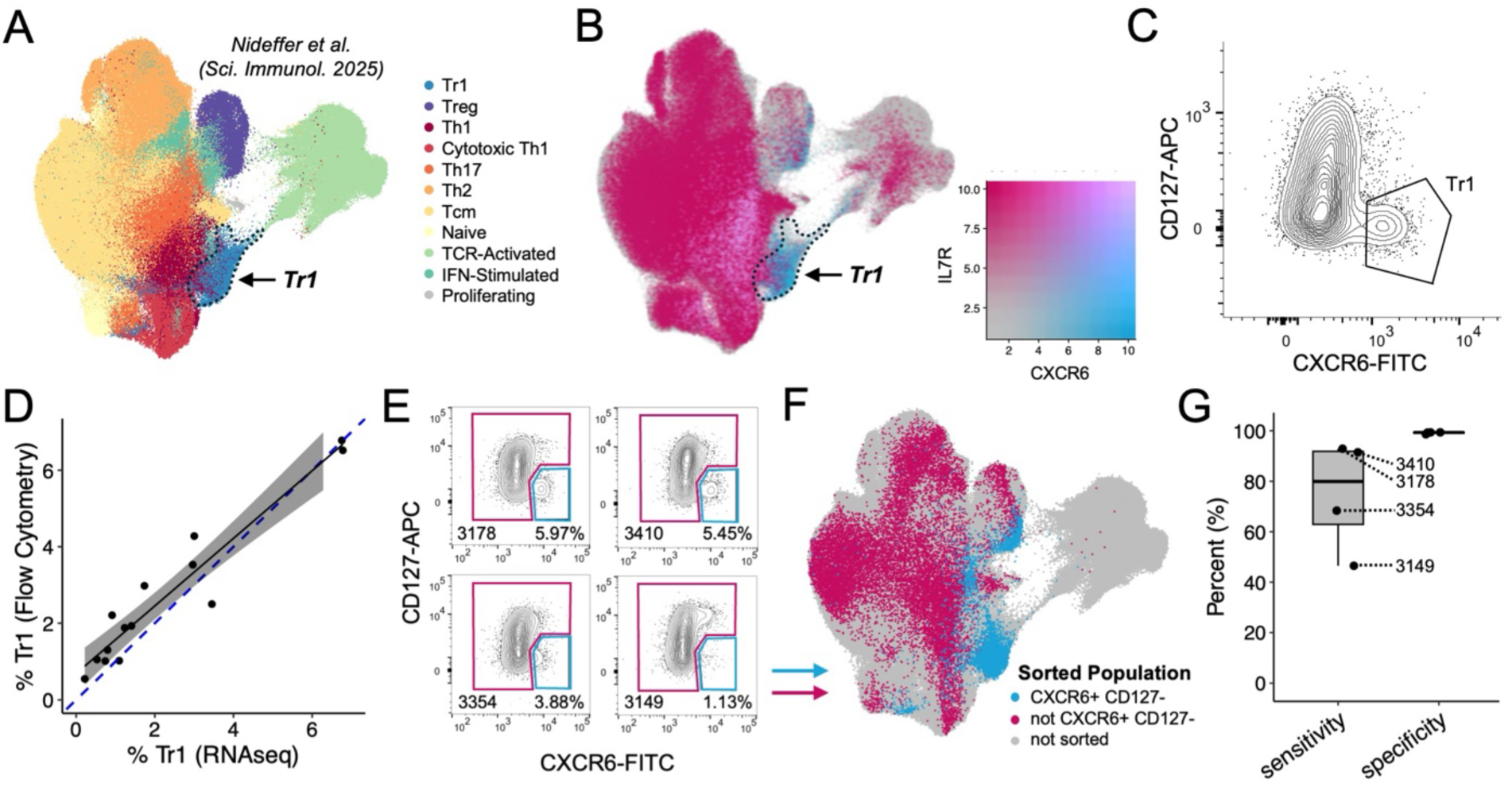
Validation of CXCR6+ CD127-as a gating strategy to enrich for Tr1 cells. (**A**) UMAP from Nideffer et al., where memory CD4+ T cells from Ugandan children and adults were subjected to scRNA/TCRseq. The dashed line highlights the resting Tr1 population, which is transcriptionally and clonally distinct from other subsets. (**B**) The same UMAP as ‘A’, but cells are colored according to their dual expression of *IL7R* and *CXCR6*. (**C**) Flow cytometry gating of memory CD4+ T cells that are CXCR6+ CD127-. (**D**) Linear relationship between the frequency of Tr1 cells determined by flow cytometry (CXCR6+ CD127-) versus scRNAseq (unsupervised clustering). A 95% confidence range for the linear regression is depicted in gray. The dashed, blue line represents y = x. (**E**) Flow cytometry plots depicting populations from four different Ugandan children that were sorted and then analyzed by scRNAseq. (**F**) Mapping of the populations sorted in ‘E’ onto the UMAP from ‘A’ based on transcriptomics. (**G**) The sensitivity and specificity of the CXCR6+ CD127-gate for identifying Tr1 cells from memory CD4+ T cells.

Other gating strategies have been proposed for identifying Tr1 cells—CD49b+ LAG-3+ as well as CCR5+ PD-1+^10,11^. However, these populations demonstrated only partial overlap with CXCR6+ CD127-Tr1 cells (Fig. 2A). Even though the CCR5+ PD-1+ and LAG-3+ CD49b+ gating strategies both enrich for IL-10-expressing cells (Fig. 2B), these gates only capture approximately 15% of the memory CD4+ T cells that make IL-10 in response to PMA/Ionomycin stimulation (Fig. 2C-D). In contrast, the CXCR6+ CD127-gating strategy captures ∼58% of IL-10-expressing memory CD4+ T cells (Fig. 2C-D). While IL-10 expression alone should not be the benchmark of good Tr1 surface markers^13^, these data are consistent with our scRNAseq validation, and support the use of CXCR6 and CD127, rather than other markers, for the approximate identification of Tr1 cells in the context of malaria.

**Fig. 2.**
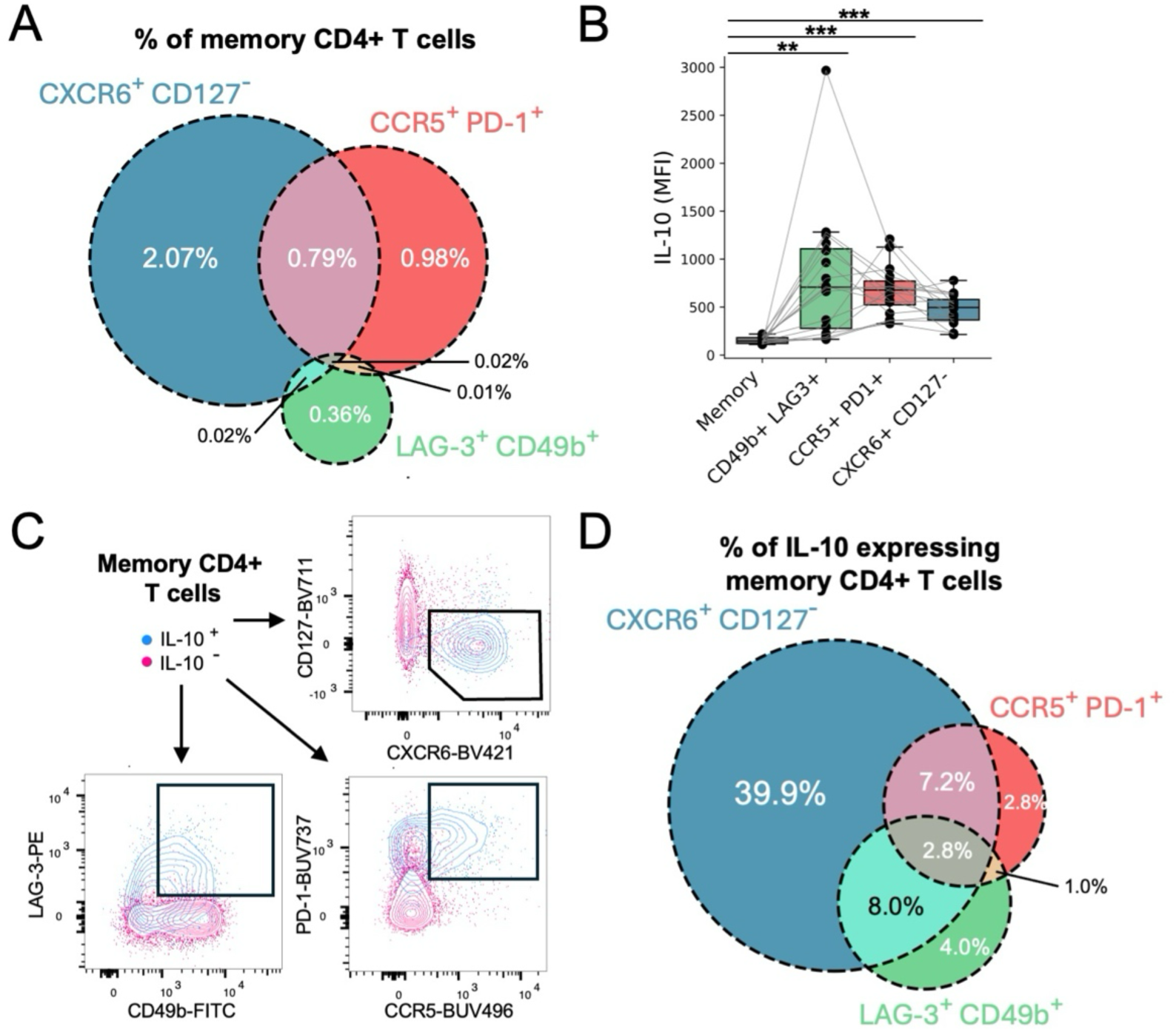
Comparison between the CXCR6+ CD127-gating strategy with previously proposed gating strategies for Tr1 cells. (**A**) Venn diagram depicting the overlap of memory CD4+ T cells that are CXCR6+ CD127-, CCR5+ PD-1+, and/or LAG-3+ CD49b+. (**B**) Mean fluorescence intensity (MFI) of IL-10 among different populations stimulated with PMA and ionomycin. Each dot represents the MFI of a population within a single donor. Lines connect populations from the same donor. Significance was determined via paired T tests with multiple hypothesis test correction. p-value < 0.01 = **; p-value < 0.001 = ***. (**C**) Flow cytometry plots depicting the patterns of surface marker expression by IL-10 expressing and non-expressing memory CD4+ T cells stimulated with PMA and ionomycin. (**D**) Venn diagram depicting the overlap in the phenotypes of cells that express IL-10 in response to PMA and ionomycin.

### Tr1 cells of Ugandan children are functionally suppressive

A core function of Tr1 cells is the ability to suppress the proliferation of other CD4+ T cells via secretion of IL-10^7,14^. We, thus, performed an *in vitro* suppression assay using Tr1 cells derived from malaria-exposed children (Fig. 3A). “Suppressor” cells—Tr1 (CXCR6+ CD127-), Treg (CD25+ CD127-), or T helper (Th) cells—were sorted from peripheral blood mononuclear cell (PBMC) samples of malaria-exposed Ugandan children and then co-cultured with allogenic “responder” memory CD4+ T cells for four days in the presence of αCD3 and αCD28 (Fig. 3A-B). Labeling of suppressors with CellTrace Violet (CTV) and responders with CellTrace CFSE enabled the quantification of proliferation following co-culture (Fig. 3C). Responder CD4+ T cells proliferated extensively in the presence of Ugandan-derived Th cells; however, their proliferation was significantly suppressed in the presence of Tr1 cells and Tregs (Fig. 3C-E). Additionally, Tr1 cells, themselves, did not proliferate extensively in co-culture (Fig. 3C,F), consistent with prior findings^15,16^. IL-10 receptor blockade partially restored responder proliferation in the presence of Tr1 cells (and to a lesser extent in the presence of Tregs) (Fig. 3G-H), demonstrating that suppression by CXCR6+ CD127-Tr1 cells is partially mediated by IL-10. Interestingly, IL-10 also seemed to signal in an autocrine fashion to inhibit Tr1 proliferation, though this observed effect was modest (Fig. S3). In sum, these data confirm that CXCR6+ CD127-Tr1 cells are functionally suppressive and may therefore influence clinical immunity to malaria.

**Fig. 3.**
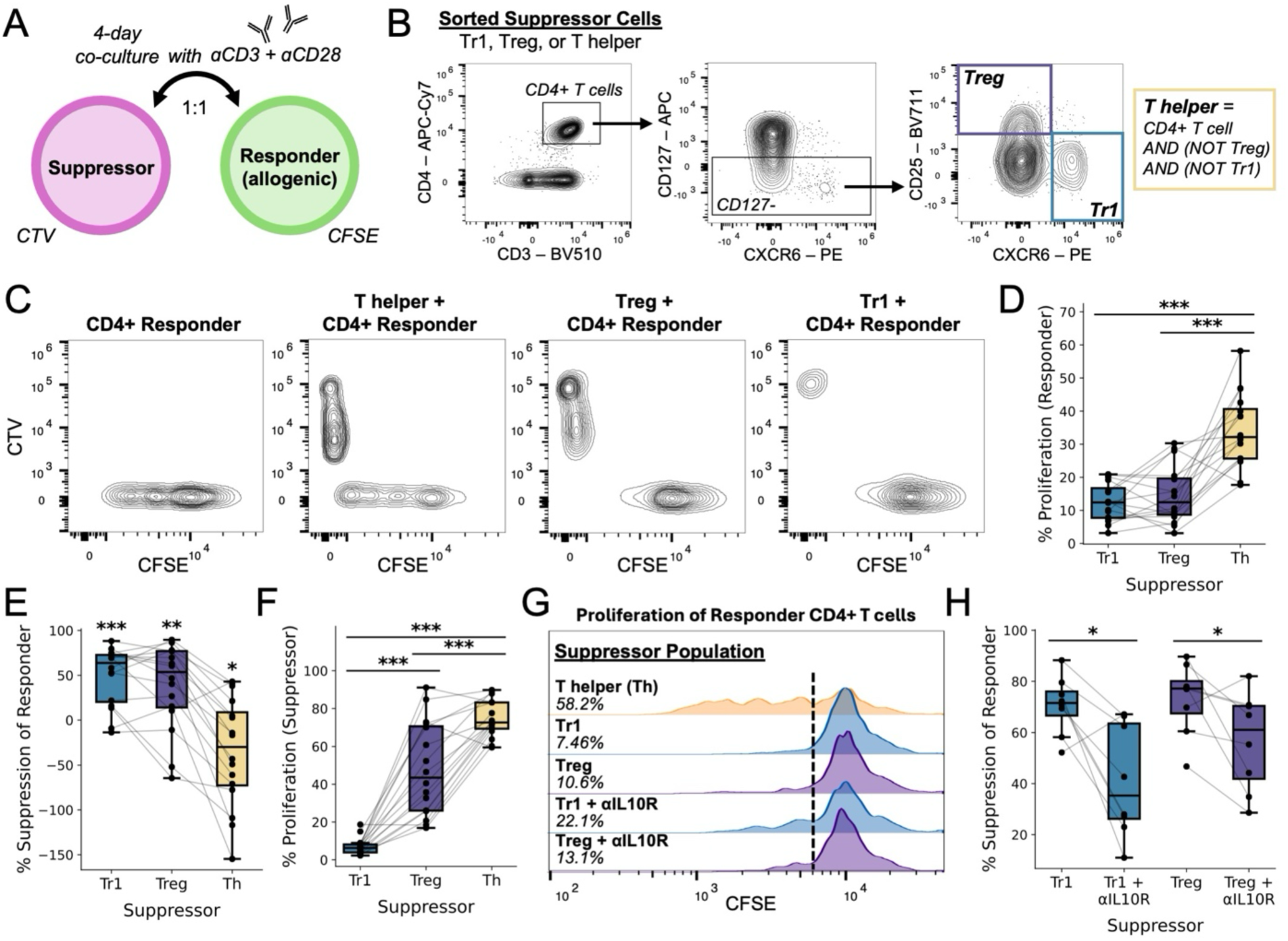
Suppression of CD4+ T cell proliferation by CXCR6+ CD127-Tr1 cells of Ugandan children. Schematic depicting experimental design, where a sorted “suppressor” population from a Ugandan child was co-cultured with allogenic “responder” memory CD4+ T cells at a ratio of 1:1. The “suppressors” and “responders” were labelled with CTV and CFSE, respectively, and co-cultured for 4 days in the presence of soluble αCD3 and αCD28 before analysis by flow cytometry. (**B**) Sorting strategy for isolating the “suppressor” populations prior to co-culture. (**C**) Flow cytometry plots after co-culture depicting the proliferation of “suppressors” and “responders”. (**D**) Percent of “responders” that are proliferating (CFSE-low) after co-culture with one of the three “suppressor” populations. Lines connect different “suppressors” derived from the same donor. (**E**) Percent suppression of the “responders” in the presence of different “suppressors.” This metric was calculated as: (“percent proliferation without suppressor” minus “percent proliferation with suppressor”) divided by “percent proliferation without suppressor.” The mean was compared to zero to determine statistical significance. (**F**) Percent of “suppressors” that are proliferating after the co-culture. (**G**) Representative histograms depicting “responder” proliferation in the presence of different “suppressors” and IL-10 receptor blockade (αIL10R) or an isotype control. Dashed line depicts the cutoff for what is deemed CFSE-low. (**H**) Percent suppression of the “responders” in the presence of either Tr1 or Treg “suppressors” with or without IL-10 receptor blockade. Significance was determined via paired T tests with correction for multiple hypotheses when appropriate. p-value < 0.05 = *; p-value < 0.01 = **; p-value < 0.001 = ***.

### CXCR6+ CD127-Tr1 cells dominate the malaria-specific response

To evaluate the clinical relevance of CXCR6+ CD127-Tr1 cells in malaria, we leveraged samples collected from individuals enrolled in the Malaria in Uganda Systems Biology and Computational Approaches (MUSICAL) Study, a longitudinal cohort study that incorporated active and passive case finding with regimented follow-up for episodes of symptomatic malaria and asymptomatic parasitemia (Table S1). We utilized peripheral blood mononuclear cells (PBMC) and plasma collected before, during, and after paired symptomatic and asymptomatic infections in the same N=48 children to study the dynamics of CD4+ T cell responses and the plasma proteome^17^ over the course of infections (Fig. 4A). The order of symptomatic and asymptomatic episodes was random for each child, and the distribution of ages for symptomatic and asymptomatic episodes were similar (Fig. 4B).

**Fig. 4.**
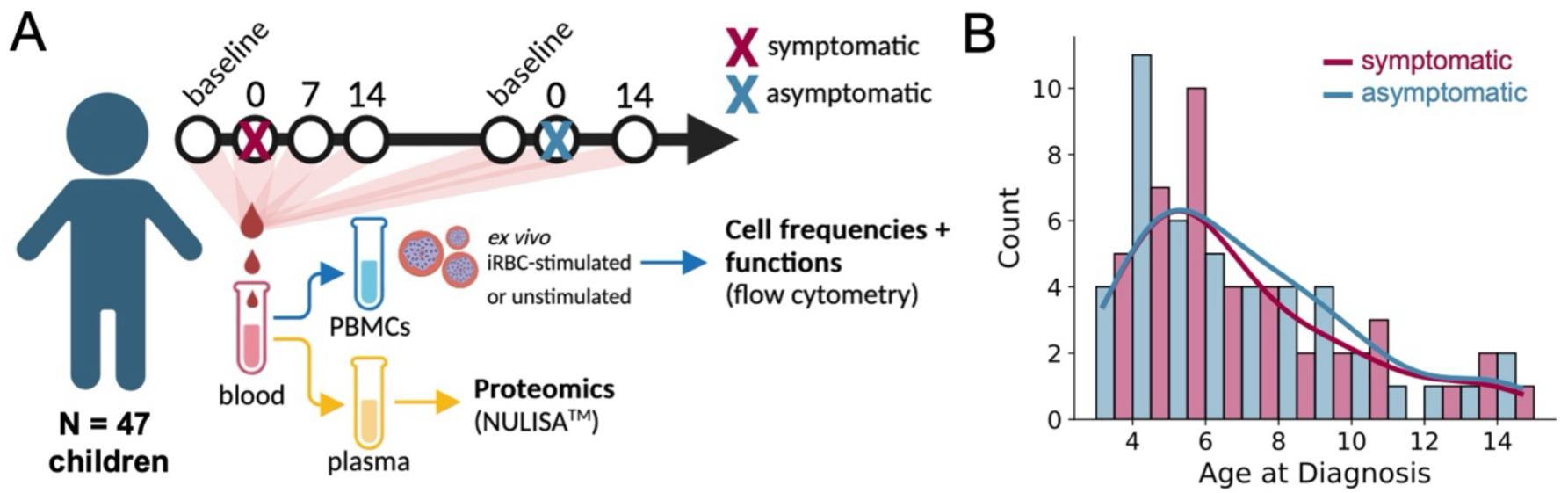
Sampling and experimental pipeline for longitudinally studying paired symptomatic and asymptomatic infections among Ugandan children and adults. (**A**) Timeline depicts how samples were provided in the context of symptomatic and asymptomatic infections experienced by the same individual. Plasma and PBMCs derived from the same blood sample were analyzed by flow cytometry and NULISA, respectively. (**B**) Histogram depicting the ages (at diagnosis) of children followed in this study. Timepoints from symptomatic and asymptomatic infections are included for each individual. Lines represent smoothed average counts.

We utilized a large multiparameter phenotyping panel to simultaneously detect Tr1 (CXCR6+ CD127-), Th1 (CCR4-CCR6-CXCR3+), Th2 (CCR4+ CCR6-CXCR3-), Th17 (CCR4+ CCR6+ CXCR3-), Treg (CD25+ FOXP3+), and circulating T follicular helper (cTfh) cells (CXCR5+ PD-1+) cells using conventional gating strategies (Fig. S4A). This panel also included key functional markers and cytokines (Fig. S4B). In response to stimulation with PMA/Ionomycin, CXCR6+ CD127-Tr1 cells were the primary producers of IL-10, though CCR4-CCR6-CXCR3+ Th1 cells also expressed some IL-10 (Fig. 5A). In response to stimulation with *Plasmodium*-infected red blood cells (iRBCs), the same pattern was observed. CXCR6+ CD127-Tr1 cells strongly upregulated IL-10 in response to iRBC stimulation, while CXCR5+ PD-1+ cTfh and CCR4-CCR6-CXCR3+ Th1 cells only modestly upregulated IL-10 (Fig. 5B). CCR4-CCR6-CXCR3+ Th1 cells were the primary producers of IFNγ in response to PMA/ionomycin (Fig. 5C). However, in response to iRBCs, CXCR6+ CD127-Tr1 cells upregulated IFNγ to the greatest extent (Fig. 5D). These data suggest that CXCR6+ CD127-Tr1 cells represent the dominant malaria blood-stage-specific CD4+ T cell population and that IFNγ during pediatric malaria is primarily Tr1 rather than Th1-derived.

**Fig. 5.**
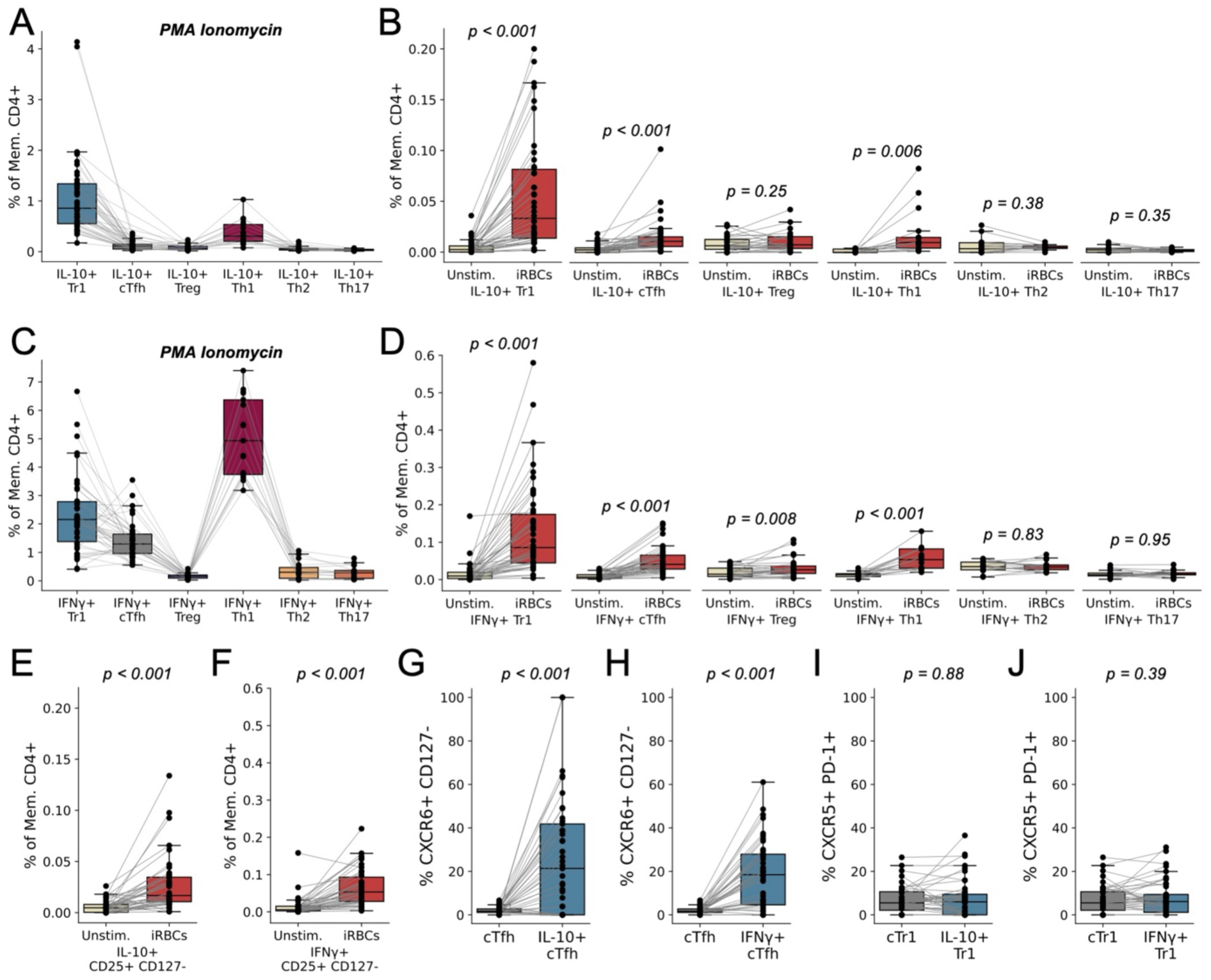
Cytokine production by memory CD4+ T cell subsets following stimulation. (**A-D**) The percentage of memory CD4+ T cells that belonged to a given subset (defined by surface markers) and produced IL-10 (‘A’, ‘B’) or IFNγ (‘C’, ‘D’) in response to stimulation with PMA and ionomycin (‘A’, ‘C’) or in response to iRBC stimulation versus unstimulated (‘B’, ‘D’). (**E-F**) The percentage of memory CD4+ T cells that were CD25+ CD127- and expressed either IL-10 (‘E’) or IFNγ (‘F’) in response to iRBC stimulation (compared to unstimulated). (**G-H**) The percentage of CXCR5+ PD-1+ cTfh and cytokine-expressing (‘G’, IL-10; ‘H’, IFNγ) CXCR5+ PD-1+ cTfh that are also CXCR6+ CD127-(stimulated with iRBCs). (**I-J**) The percentage of CXCR6+ CD127-Tr1 cells and cytokine-expressing (‘I’, IL-10; ‘J’, IFNγ) CXCR6+ CD127-Tr1 cells that are also CXCR5+ PD-1+. Significance was determined via paired T tests; p-values are displayed for all tested comparisons.

CD25+ FOXP3+ Tregs did not appreciably upregulate IL-10 or IFNγ in response to iRBC stimulation (Fig. 5B,D), but CD4+ T cells that were CD25+ CD127-(often assumed to be conventional Tregs) did respond to iRBC stimulation by upregulating IL-10 and IFNγ (Fig. 5E,F). Given our prior transcriptional data^6^, it is likely that CD25+ CD127-cells represent activated Tr1 cells, especially following *Pf* antigen stimulation. Furthermore, although CXCR5+ PD-1+ cTfh upregulated IL-10 and IFNγ upon stimulation with *Pf* antigen, the response was much lower than CXCR6+ CD127-Tr1 cells (Fig. 5B,D). There was minimal overlap observed between the larger CXCR5+ PD-1+ cTfh and CXCR6+ CD127-Tr1 populations, but nearly 20% of cytokine-producing, *Pf*-reactive cTfh could also be classified as CXCR6+ CD127-Tr1 cells (Fig. 5G,H). Similar enrichment for cTfh markers amongst *Pf*-responding Tr1 cells was not observed (Fig. 5I,J), suggesting that CXCR6 and CD127 better enrich for *Pf*-specific CD4+ T cells than CXCR5 and PD-1. Moreover, CXCR6+ CD127-Tr1 cells, to a greater extent than CXCR5+ PD-1+ cTfh, upregulated IL-21 in response to iRBCs (Fig. S5), consistent with our prior transcriptional data^6^.

### Tr1 cells and their effector molecules increase during malaria

Next, we leveraged the longitudinal aspect of MUSICAL to study CD4+ T cell subset dynamics over the course of symptomatic and asymptomatic infections. Among children, symptomatic malaria induced a rapid, ∼1.6-fold expansion in the frequency of CXCR6+ CD127-Tr1 cells but had no significant effect on other CD4+ T cell subsets (Fig. 6A), consistent with our prior findings using scRNA/TCR seq^6^. This trend was similarly observed in adults who contracted malaria, though our sample size was limited to 4 individuals (Fig. S6). Because malaria causes lymphopenia (Fig. S7), the absolute number of Tr1 cells in peripheral blood was not elevated during acute malaria (even though relative Tr1 frequencies were elevated) (Fig. 6B). Asymptomatic infection did not significantly affect the frequency or absolute number of Tr1 cells, although a trending increase was observed (Fig. S8). The frequency of Tr1 cells that expressed IL-10, IFNγ, and PD-1 increased in both symptomatic malaria and asymptomatic parasitemia (Fig. 6C-E). Concurrently, some Tr1 cells upregulated their surface expression of CXCR5 (Fig. S9), suggesting that some Tr1 cells may traffic to secondary lymphoid organs.

**Fig. 6.**
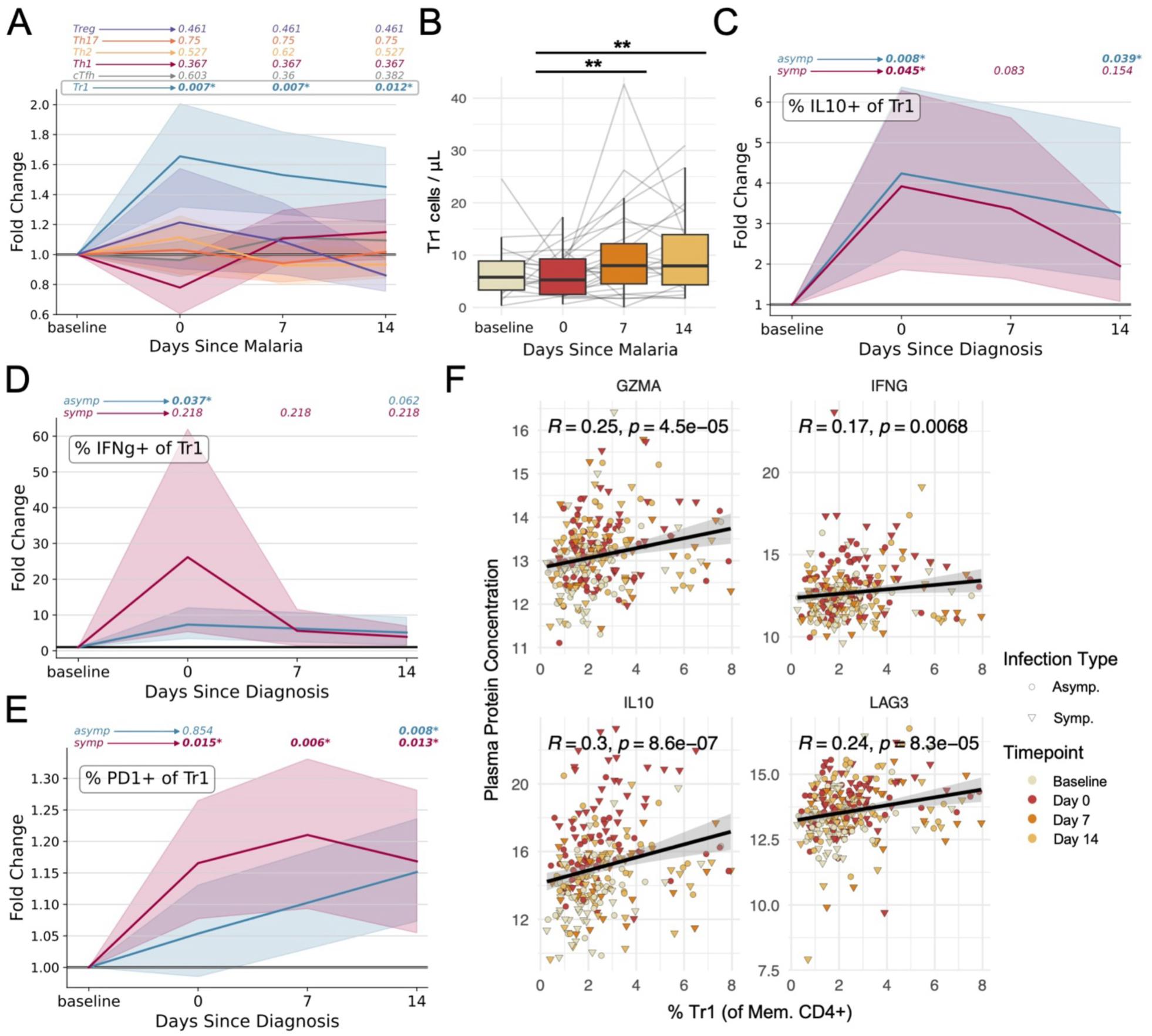
Infection dynamics of memory CD4+ T cells populations and effector molecules. (**A**) Fold change (compared to the pre-infection baseline) in cell frequencies following symptomatic malaria as determined by flow cytometry. (**B**) The absolute counts of Tr1 cells in peripheral blood before, during, and after symptomatic malaria. (**C-E**) Fold change in the percentage of Tr1 cells expressing IL-10 (‘C’), IFNy (‘D’), or PD-1 (‘E’) following symptomatic malaria or asymptomatic parasitemia. For fold change plots, p-values are displayed above each sample timepoint for each population and represent pair-wise comparisons to baseline. (**F**) Correlations between plasma concentrations of granzyme A, IFNγ, IL-10, and LAG3 (determined by NULISA) and Tr1 frequencies (determined by flow cytometry). The color and shape of individual points denotes the timepoint of the sample and whether the infection was symptomatic or asymptomatic. Gray area represents 95% confidence range. Pearson’s R and associated p-values are displayed above each plot. Unless the p-value is explicitly reported, p-value < 0.05 = *; p-value < 0.01 = **; p-value < 0.001 = ***

To understand the influence of Tr1 cells on the systemic response to infection, we quantified the abundance of Tr1-associated molecules (granzyme A, IFNγ, IL-10, and LAG3) in plasma from the same blood samples used to study CD4+ T cells as a part of MUSICAL. The genes encoding granzyme A, IFNγ, IL-10, and LAG3 were all associated with resting and/or activated Tr1 cells (Fig. S10). Plasma concentrations of all four of these proteins correlated with Tr1 cell frequencies (Fig. 6F), suggesting that Tr1 cells likely influence the systemic levels of these molecules. Furthermore, the plasma concentrations of these Tr1 molecules increased during symptomatic malaria (Fig. S11). This is perhaps not surprising, since plasma concentrations of 112 proteins (out of 250 proteins measured) were differentially abundant during malaria (Fig. S12A). However, in the case of asymptomatic parasitemia, only 1 protein was significantly elevated at the time of diagnosis—IL-10 (Fig. S12B). Thus, asymptomatic parasitemia was distinct from symptomatic malaria in that a canonical Tr1 effector molecule but not inflammatory mediators were elevated in plasma at diagnosis. Interestingly, plasma concentrations of granzyme A, IFNγ, IL-10, and LAG3 were higher on day 14 after asymptomatic infection among individuals that were still parasitemic compared to those that had cleared their infection by day 14 (Fig. S13), suggesting a link between persistent parasitemia and persistent Tr1 responses. Altogether, these data suggest that Tr1 cells affect the systemic immune response to *Plasmodium falciparum,* and that this may potentially influence clinical outcomes of infection.

### Tr1 cell influence clinical immunity and parasite persistence

Given the dominance of Tr1 cells in pediatric malaria and their strong regulatory potential in other contexts, we hypothesize that these cells might influence malaria disease outcomes. We tested whether Tr1 cell frequencies prior to diagnosis were associated with the probability of symptoms given infection. Models were adjusted for age as Tr1 cell frequencies positively correlated with this variable (Fig. S14). Higher frequencies of Tr1 cells prior to infection correlated with a significantly lower probability of developing symptoms upon infection when accounting for parasitemia and individual variability (Fig. 7A). Every 1% increase in Tr1 frequencies was associated with a ∼3.2-fold decrease in the odds of symptoms given infection, and an ability to tolerate a ∼3.0-fold increase in parasite density (Fig. 7A-B). We did not observe a significant relationship between Tr1 cell frequencies measured either prior to (coef. = 0.02; 95% CI: - 0.11 - 0.15) or at the time of infection (coef. = 0.11; 95% CI: - 0.02 - 0.23) and log10 parasite densities at the time of infection.

**Fig. 7.**
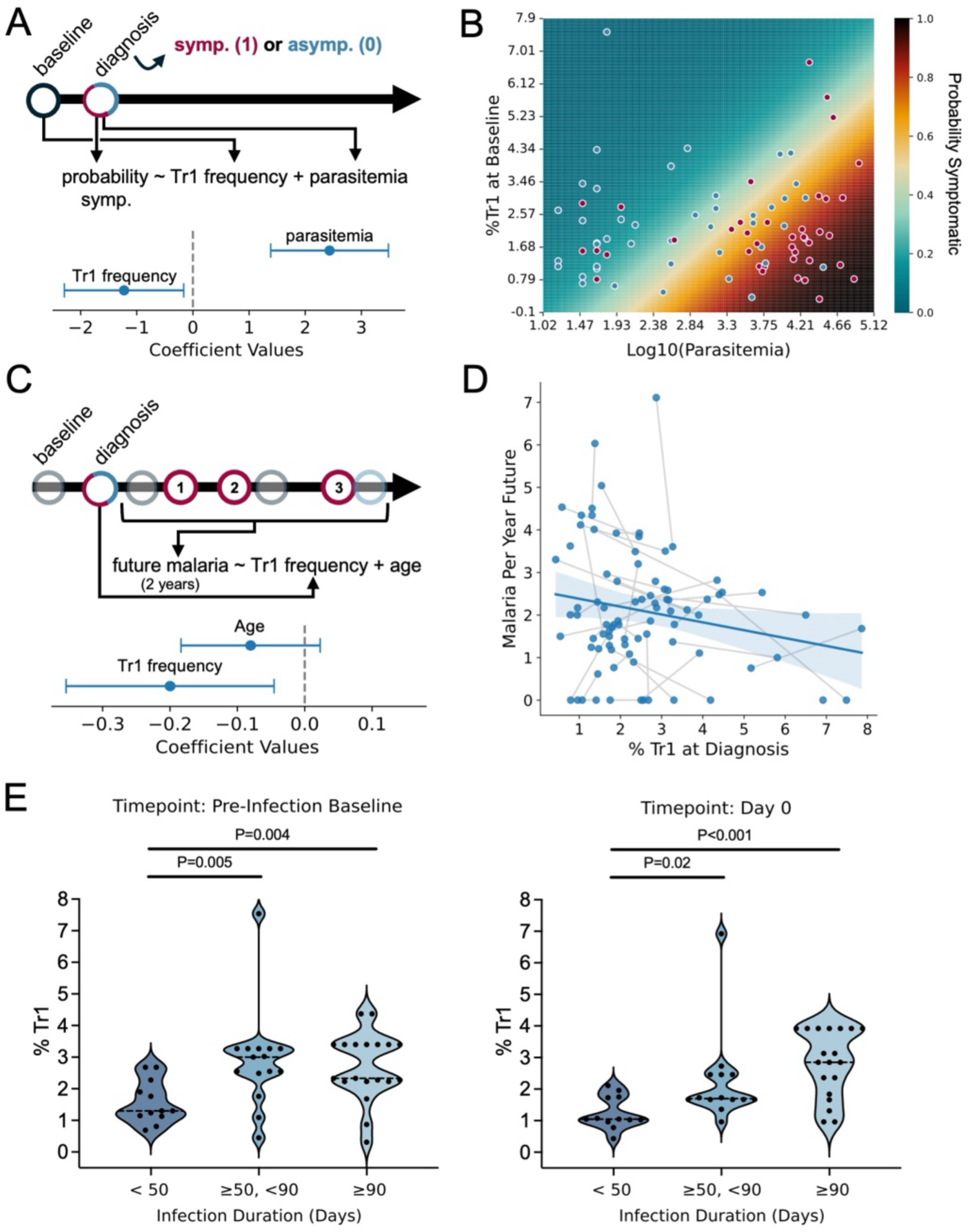
Correlations between Tr1 cells and outcomes of *Pf* infections. (**A**) Schematic describing the logistic regression model to predict whether an infection will be symptomatic (1) or asymptomatic (0) based on Tr1 cell frequencies at the pre-infection baseline timepoint and log-10 transformed parasite density at the time of diagnosis. Fitted coefficients are displayed with 95% confidence intervals below the schematic (**B**) The fitted probability (model from ‘A’) that an infection will be symptomatic based on various combinations of parasitemia at diagnosis and Tr1 frequencies prior to diagnosis. Dots represent true observations used to fit the model (red indicates symptomatic; blue indicates asymptomatic). (**C**) Schematic describing the generalized linear model to predict the 2-year future incidence of malaria based on age and Tr1 frequencies at diagnosis. Fitted coefficients are displayed with 95% confidence intervals below the schematic. (**D**) Scatterplot depicting the relationship between Tr1 frequencies at diagnosis and the future incidence of malaria. Gray lines connect two separate observations from the same individual. Blue line and shaded 95% confidence area depict a simple linear regression between %Tr1 at diagnosis and malaria incidence per person year (does not account for repeated measures or age.) (**E**) %Tr1 measured at the pre-infection timepoint (left) and at diagnosis (right) stratified by duration of incident, untreated asymptomatic infections (as determined by AMA1 amplicon sequencing). Groups compared by non-parametric Wilcoxon Rank-sum test. p-value < 0.05 = *; p-value < 0.01 = **; p-value < 0.001 = ***.

While these data suggest that the frequency of Tr1 cells influences the probability of symptoms given the next, immediate infection, we additionally hypothesized that the response to a single infection might influence future outcomes. Thus, we evaluated whether Tr1 frequencies measured during infection were associated with the incidence of subsequent symptomatic malaria (Fig. 7C), accounting for repeated measures within individuals (multiple infections) and controlling for age. Tr1 frequencies at the time of infection were associated with a lower incidence of symptomatic malaria—where a 5% increase in Tr1 cell frequency was predicted to reduce malaria incidence by ∼1 episode per year (Fig. 7C,D; Table S2).

Finally, while Tr1 cells appeared to be associated with asymptomatic outcomes, we asked whether they might also impact parasite clearance. To test this hypothesis, we utilized longitudinal parasite genotyping by AMA1 amplicon sequencing to categorize the duration of asymptomatic infections. Then, we assessed whether infection duration was associated with Tr1 frequencies. Higher Tr1 frequencies measured prior to or at the time of diagnosis positively correlated with infection duration (Fig. 7E), independent of age, parasitemia at the time of infection, and clustering within individuals (Table S2). Together, these data support the hypothesis that Tr1 cells help mediate clinical immunity to malaria, but that Tr1-mediated disease tolerance could contribute to delayed pathogen clearance.

## Discussion

In this study, we demonstrated that CXCR6 and CD127 can be used to enrich for Tr1 cells in the peripheral blood of malaria-exposed Ugandan children. These cells strongly suppressed CD4+ T cell proliferation in a manner that was partially dependent on IL-10. We also showed that CXCR6+ CD127-Tr1 cells responded to *in vitro* stimulation with *Plasmodium* antigens by expressing IL-10, IFNγ, and IL-21 at the protein level, demonstrating their specificity for *Pf* antigens. Furthermore, Tr1 cells expanded rapidly and produced IL-10 and IFNγ *in vivo* following malaria infection. Tr1 cell frequencies correlated with concentrations of soluble granzyme A, IFNγ, IL-10, and LAG-3 in the blood, suggesting their ability to influence immune responses systemically. Importantly, the abundance of CXCR6+ CD127-Tr1 cells before and during infection correlated with future clinical outcomes, including the probability of symptoms given parasitemia and the duration of asymptomatic parasitemia. Altogether, these data highlight the clinical relevance of CXCR6+ CD127-Tr1 cells as a functionally suppressive cell subset that likely aids in the acquisition of clinical immunity to malaria but may facilitate parasite persistence.

To identify surface markers for a specific T cell subset, one must first define a reliable, biologically meaningful “ground truth” against which to validate the proposed markers. The original markers proposed for Tr1 cells—CD49b and LAG-3—were identified by gene expression profiling of IL-10–expressing, suppressive CD4+ T cells^10^. With the availability of newer technologies like single-cell RNA and TCR-seq, we can now redefine Tr1 cells based on global transcriptional profiles. In our recent study, Tr1 cells were identified as a transcriptionally and clonally distinct subset that responded uniquely to TCR stimulation and persisted with stable gene expression over time^6^. Using this higher-resolution definition, we evaluated surface markers and found that CXCR6+ CD127− cells effectively enriched for Tr1 cells in malaria-exposed children. Although LAG-3 was expressed at the RNA level and plasma levels of soluble LAG-3 correlated with Tr1 abundance, it was not an effective surface marker—possibly due to shedding or intracellular localization, consistent with prior studies showing proteolytic cleavage of LAG-3 from CD4+ T cells^18^. Importantly, neither CXCR6 and CD127 nor any other surface marker combination should be viewed as definitive markers of Tr1 identity, but as practical tools to enrich for cells that are more precisely defined by a complex and stable transcriptional program, exhibiting clonal fidelity. As immunology moves toward more reproducible and scalable cell classification systems, high-parameter methods such as single-cell RNA and ATAC sequencing will be essential for defining T cell ontologies, and surface markers for all subsets should be validated by sequencing sorted populations, as done here.

We utilized these newly validated surface markers to study the CD4+ T cell response to *P*. *falciparum*. We found that CXCR6+ CD127-Tr1 cells represent the dominant malaria blood-stage-specific CD4+ T cell population, consistent with our prior scRNAseq/TCRseq study^6^. CXCR6+ CD127-Tr1 cells respond to *Plasmodium* antigen stimulation *in vitro* and *in vivo* by upregulating IL-10 and IFNγ, and they expand following symptomatic disease, more so than any other cellular subset, including CCR4-CCR6-CXCR3+ Th1 cells, CXCR5+ PD1+ cTfh cells, or CD25+ CD127-Tregs. Despite this, prior studies have described the expansion of CXCR5+ PD-1+ cTfh following *Plasmodium* infection^19–21^. In our study, we did observed CXCR5+ PD-1+ cells that produced IL-10 and IFNγ in response to iRBCs (Fig. 5B,D). However, many of these cells were also CXCR6+ CD127-(Fig. 5G,H), suggesting that marker-based definitions alone may not adequately discriminate between these different cellular populations. Future studies should employ single-cell sequencing of sorted CXCR5+ PD-1+ CD4+ T cells to determine if these cells represent a distinct but transcriptionally unified population or if the CXCR5+ PD-1+ population is heterogeneous and comprises cells from each of the various helper and regulatory subsets.

After establishing that CXCR6+ CD127-Tr1 cell were the predominant CD4+ T cell population responding to malaria in children, we leveraged the longitudinal, paired nature of the MUSICAL clinical study to determine whether these cells correlated with clinical outcomes of infection. Clinical immunity to malaria is thought to be mediated by immunological mechanisms balancing disease tolerance (i.e., probability of symptoms given parasitemia) and parasite control (i.e. parasite densities given infection)^22^. We do not expect that Tr1 cells are solely responsible for clinical immunity—they are a regulatory subset that may support anti-disease immunity, but they may not contribute to anti-parasite immunity. Indeed, we did not observe a significant correlation between Tr1 frequencies and parasite densities at the time of infection. In contrast, we found a significant correlation between frequencies of Tr1 cells and the probability of symptoms given parasitemia. Our analysis provides evidence that Tr1 cells contribute to anti-disease immunity but that other mechanisms controlling parasite growth, such as antibodies, B cells^23^, γδ T cells^24^, and natural killer cells^25^, are also likely important for protection following repeated infections.

Although Tr1 cells correlated with anti-disease immunity, higher frequencies of these cells were also associated with delayed parasite clearance among children with asymptomatic, untreated infections. Antigen-specific Tr1 cells have long been hypothesized to contribute to chronic infection and/or susceptibility to re-infections^8^. In murine models, IFNã- and IL-10-producing CD4^+^ T cells emerge during experimental infection with *Toxoplasma gondii* and in non-healing cutaneous leishmaniasis, where they were shown to protect mice against immune-related pathology at the cost of pathogen persistence^26,27^. In another murine model of leishmaniasis, CD4+ CD25+ cells accumulated at the site of infection, regulated the function of local effector cells, and prevented efficient elimination of the parasite^28^. In a murine model of *Plasmodium yoelii,* depletion of CD4+ CD25+ Tregs protected mice from death by restoring an effector immune response that efficiently eliminates parasites^29^. Nevertheless, evidence supporting a role for Tr1 cells in interfering with pathogen clearance in humans has been sparse^8^. The unique nature of the MUSICAL longitudinal cohort, with individual, genotyped infections, enabled careful consideration of infection duration.

Although our study suggests that Tr1 cells correlate with asymptomatic outcomes, these cells did not expand as prolifically following asymptomatic parasitemia compared to symptomatic malaria. We further show that Tr1 cells are highly suppressive but also quite anergic relative to other CD4+ T cell subsets (Fig. 3D-F). Therefore, memory Tr1 cells that are activated upon reinfection may be slow to proliferate. This could explain why, during asymptomatic infection, preexisting Tr1 cells respond to antigen and produce cytokines *in vivo* but do not proliferate extensively (Fig. 6C,D, Fig. S8). Furthermore, if there are sufficient preexisting, suppressive Tr1 cells to adequately influence immune responses and clinical outcomes to a new infection, it would be expected that CD4+ T cell expansion would be strongly suppressed. It may be the case that symptomatic malaria results in part from the inability of preexisting *Pf-*specific Tr1 memory cells to recognize the antigens of a newly infecting strain. In this setting, Tr1 expansion might be driven by the activation of naïve CD4+ T cells, which may proliferate more readily compared to the somewhat anergic, memory Tr1 cells.

There are limitations to this study. We identified CXCR6 and CD127 as effective markers for enriching for Tr1 cells in a specific tissue and clinical context—the blood of malaria-exposed humans. It is, thus, uncertain whether these markers can be applied universally to enrich for Tr1 cells in other contexts. Future studies to enrich for Tr1 cells in a different clinical context should validate these markers (ideally using cell sorting and single-cell sequencing). We also correlated Tr1 frequencies with clinical outcomes, but presumably, not all Tr1 cells of Ugandan children are *Pf*-specific (though many of them are). Furthermore, *Pf*-specific Tr1 clones likely recognize diverse antigens and may not respond to the same *Pf* strains. Therefore, while Tr1 abundance prior to an infection seems to be important for clinical outcomes, it is likely that the specificity of preexisting Tr1 memory cells is also highly relevant. This could explain certain instances in which a child had many Tr1 cells prior to infection but still contracted symptomatic malaria. To this end, it is important that future studies explore the specific *Pf* peptides that are recognized by Tr1 cells. Finally, functional experiments assessed the capacity of Tr1 cells to suppress allogeneic T cell proliferation. Although, this form of suppression is likely an important functional feature of Tr1 cells, future experiments will need to identify whether Tr1 cells have other functions (e.g., inhibition of the inflammasome or killing of antigen presenting myeloid cells) as identified in other contexts^9,30,31^.

By validating new cell surface markers (CXCR6+ CD127-) using scRNAseq and functional experiments to identify high-confidence, bona-fide Tr1 cells, we were able to study the function and clinical relevance of Tr1 cells among children living in malaria-endemic Uganda. Our data demonstrate that CXCR6+ CD127-Tr1 dominate the response to *Pf* infected red blood cell antigens, rapidly expand following malaria *in vivo*, and express IL-10 and IFNγ *in vivo* during both symptomatic malaria and asymptomatic parasitemia. Further, these cells may mediate both the tolerance and persistence of parasitemia through IL-10 and other mechanisms of immune regulation. This work updates prior notions of what defines a Tr1 cell and provides important insights into a potential mediator of clinical immunity to malaria.

## Materials and methods

### Clinical studies and sample collection

This work utilizes samples provided by Ugandan children and adults living in Tororo or Busia who were part of the East African International Centres of Excellence in Malaria Research (ICEMR) cohorts. In this setting, malaria transmission is year-round, with two seasonal peaks. Informed consent was provided by all participants or by their parents/legal guardians. Screening and enrollment of households in the parent ICEMR cohort have been previously described^32,33^. Briefly, households in a contiguous study area across Busia and Tororo districts were enumerated to provide a sampling frame, and 80 households were randomly selected for screening and enrolled if they met the following criteria: 1) having at least two members aged 5 years or younger; 2) no more than 7 permanent residents currently residing; 3) no plans to move from the study catchment area in the next 2 years; and 4) willingness to participate in entomological surveillance studies. Cohort participants were followed for all care at dedicated study clinics. Participants were monitored for parasitemia by microscopy and qPCR by active (monthly) and passive surveillance, with peripheral blood samples taken. Those who presented with a fever (tympanic temperature >38.0 °C) or history of fever in the previous 24 hours had blood obtained by finger prick for a thick smear. If the thick smear was positive for *Plasmodium* parasites, the patient was diagnosed with malaria regardless of parasite density and was treated with artemether-lumefantrine. Participants with asymptomatic parasitemia were not treated with antimalarial drugs in accordance with local guidelines. In the MUSICAL subcohort, study participants diagnosed with symptomatic malaria were asked to return for repeat clinic visits and blood sampling on days 7, 14, and 28, post-treatment. Those with asymptomatic parasitemia were asked to return for repeat clinic visits and blood sampling on days 14 and 28 following diagnosis (they were not treated). PBMCs were isolated from peripheral blood samples by density gradient centrifugation; these samples were then cryopreserved and shipped by liquid nitrogen to Stanford University for further analysis.

The study protocols were approved by the Uganda National Council of Science and Technology, the Makerere University School of Medicine Research and Ethics Committee, the University of California, San Francisco Committee on Human Research, and the Institutional Review Board at Stanford University.

### Tr1 marker validation using scRNA/TCRseq

Cryopreserved PBMC samples from four different Ugandan children were thawed and then stained with antibodies: from BioLegend, anti-CXCR6-FITC (K041E5), anti-CD127-APC (A019D5), anti-CD3-BV421 (SK7), anti-CD4-BV785 (SK3); from BD, anti-CD8a-BV605 (SK1). Cell hashing was also performed prior to sorting so that cells from each of the four donors could be identified via distinct molecular barcodes after pooling samples together for single-cell capture (using BioLegend TotalSeq^TM^-C0251, C0252, C0253, and C0254). Tr1 cells (CXCR6+ CD127-) were pooled together and loaded into a single well on a Chromium X, and the same was done for “non-Tr1” cells (not CXCR6+ CD127-). After single-cell capture, preparation of RNA, TCR, and hashtag libraries was performed according to 10X Genomics published protocol (CG000330: Chromium Next GEM Single Cell 5-v2 Cell Surface Protein UserGuide RevD). Paired-end sequencing of the prepared libraries was performed using the NovaSeq X Plus, followed by demultiplexing of pooled libraries.

Processing of sequencing data was performed as previously described^6^ with raw FASTQ files being used to generate filtered feature-barcode matrices and TCR annotations following Cell Ranger Count v7.1.0 (reference Human GRCh38 2020-A) and Cell Ranger V(D)J v7.1.0. Further data processing (including filtering low-quality cells, annotating, clustering, performing UMAP) was performed using Seurat as previously described^6^. Finally, the sequenced cells were analyzed for their purity to validate CXCR6+ CD127-as a viable gating strategy to enrich for Tr1 cells.

### Flow cytometric analyses

Cryopreserved PBMC samples were thawed, counted, and resuspended in R10 (RPMI 1640 supplemented with 50 U/mL of penicillin, 50μg/mL of streptomycin, 2mM of L-glutamine, 10mM of HEPES, and 10% heat-inactivated fetal bovine serum). Each sample was split into three fractions subjected to different stimulation conditions: (1) unstimulated for 24 hours (250,000 PBMCs); (2) PMA/ionomycin-stimulated for 4 hours (after a 20-hour rest period) (250,000 PBMCs); (3) iRBC-stimulated for 24 hours (1,000,000 PBMCs). All samples/fractions were incubated in 200 μL of R10 in individual wells of a round-bottom 96-well plate.

*Plasmodium falciparum* blood-stage *3D7* parasites were grown as described in our prior study^6^. Once synchronous, high-density cultures were obtained, schizonts/late-stage trophozoites were purified by magnetic separation and cryopreserved prior to use in stimulation assays. For the iRBC-stimulated fractions, these aliquots were thawed, counted, and then incubated with PBMCs at a ratio of 1:2 (iRBCs:PBMCs). After 6 hours of rest/stimulation, Brefeldin A and monensin (BD Pharmingen) were added (10 μg/mL) to the unstimulated and iRBC-stimulated fractions to sequester cytokines for later staining. For the PMA/Ionomycin-stimulated fraction, Brefeldin A and monensin were added at the time of stimulation.

After 24 hours, cells were washed and stained with Live/Dead Fixable Near-IR (Thermo Scientific #L34994). Cells were again washed; the supernatant was aspirated; and then, 1 μL of each of the chemokine receptor-targeting antibodies was added to the pellets (resuspended in remaining volume after aspiration). Chemokine receptor-targeting antibodies included: from BioLegend, anti-CXCR6-BV421 (K041E5) or anti-CXCR6-FITC (K041E5), anti-CXCR3-FITC (G025H7) or anti-CXCR3-BV421 (G025H7), anti-CCR4-PE (L291H4); from BD, anti-CXCR5-BUV563 (RF8B2), anti-CCR6-BUV496 (11A9). After 15 minutes at 37°C, a cocktail of the remaining surface antibodies was added (without washing): from BioLegend, anti-CD45RA-BV785 (HI100), anti-CD127-BV711 (A019D5), anti-CD8a-BV650 (SK1), anti-CD3-AF700 (OKT3); from BD, anti-PD-1-BUV737 (EH12.1), anti-CD4-BUV395 (SK3), anti-CD25-BV605 (2A3). This cocktail was made up in equal parts PBS and BD Horizon^TM^ Brilliant Stain Buffer. Surface staining occurred in ∼50 μL for 30 minutes at room temperature in the dark. After incubation, the cells were washed in PBS, and then fixation/permeabilization was performed using the EBioscience^TM^ Foxp3/Transcription Factor Staining Buffer Set according to the manufacturer’s instructions. Then intracellular staining was performed with the following antibodies: from BioLegend, anti-IL-10-PE-Dazzle594 (JES3-19F1); from BD, anti-FOXP3-BB700 (236A/E7); from BD Pharmingen, anti-IFNγ-PE-Cy7 (B27); from eBioscience, anti-IL-21-APC (3A3-N2). After staining, the cells were washed twice more with FACS buffer (PBS with 2mM EDTA and 5mg/mL bovine serum albumin), resuspended in 150 μL, and then analyzed using a BD FACSymphony^TM^ A5 Cell Analyzer.

In certain instances, antibodies in the core panel described above were substituted for the following: from BioLegend, anti-LAG-3-PE (11C3C65); from BD, anti-CTLA-4-BB700 (BNI3), anti-CCR5-BUV496 (3A9); from eBioscience, anti-CD49b-FITC (Y418).

### Proteomic analysis using the NULISA^TM^ platform

Frozen human plasma samples were used to quantify the relative abundance of 250 circulating proteins using the NULISA^TM^ (NUcleic acid Linked Immuno-Sandwich Assay) platform, a high-sensitivity, multiplexed immunoassay. Plasma was thawed and aliquoted on ice. Individual aliquots were processed according to the manufacturer’s guidelines (NULISA, Alamar Biosciences). Briefly, each target protein was captured using a pair of matched antibodies, each conjugated to unique oligonucleotides. Upon sandwich complex formation, the proximity of the oligo-conjugated antibodies allowed for a DNA ligation reaction that generated a unique barcode sequence corresponding to each target protein.

Ligation products were then amplified by PCR and quantified via next-generation sequencing (NGS), yielding digital counts proportional to the concentration of each protein. Data were normalized using internal spike-in controls and batch correction procedures provided by the manufacturer. All samples were run in duplicate, and quality control metrics included assessment of ligation efficiency, amplification bias, and sequencing depth. Protein counts were log-transformed and scaled prior to statistical analysis.

### qPCR and parasite genotyping

For qPCR and genotyping, we collected 200 µL of blood at each visit. DNA was extracted using the PureLink Genomic DNA Mini Kit (Invitrogen) and parasitemia was quantified using an ultrasensitive varATS qPCR assay with a lower limit of detection of 0.05 parasites/µL (<u>Hofmann</u> <u>et al., 2015</u>). Samples with a parasite density >= 1 parasites/µL blood were genotyped via amplicon deep-sequencing. Hemi-nested PCR was used to amplify a 236 base-pair segment of apical membrane antigen 1 (AMA-1) using a published protocol with modifications^34^. Samples were amplified in duplicate, indexed, pooled, and purified by bead cleaning. Sequencing was performed on an Illumina NextSeq and NovaSeq platforms (150 bp paired-end). Data extraction, processing, and haplotype clustering were performed using SeekDeep^35^, followed by additional filtering^36^.

A clone was defined as a genetically identical group of parasites (e.g., with the same AMA-1 haplotype). Each unique clone was counted as an infection, and its disappearance marked a clearance event. Baseline infections were clones detected within the first 60 days of observation. Any new clone detected after day 60 was considered a new infection. To account for intermittent detection due to low parasite density or technical limitations, up to three missed detections (“skips”) were allowed before declaring a clone cleared. An infection was considered cleared only if the clone was absent in four consecutive routine samples, with the last detection date recorded as the end date. If the same clone reappeared in a subjec after being absent for at least four consecutive routine visits, it was classified as a new infection.

### Suppression assay

Cryopreserved PBMC samples from Ugandan children and one unexposed North American donor (provided through the Stanford Blood Center) were thawed. Memory CD4+ T cells from the North American donor were isolated by MACS (STEMCELL Technologies #19157) and served as an allogenic responder population. This responder population was labelled with CellTrace^TM^ CFSE (Thermo Scientific) according to the manufacturer’s instructions. Similarly, the thawed PBMC samples of Ugandan children were labelled with CellTrace^TM^ Violet (Thermo Scientific), after which, they were stained with fluorescently labelled antibodies: from BioLegend, anti-CD3-BV510 (OKT3), anti-CD25-BV711 (M-A251), anti-CXCR6-PE (K041E5), anti-CD127-APC (A019D5), anti-CD4-APC/Cy7 (RPA-T4). A Sony SH800S Cell Sorter was then used to isolate three “suppressor” populations from each Ugandan sample: (1) Tr1 cells (CD3+, CD4+, CD127-, CXCR6+, CD25-), (2) Tregs (CD3+, CD4+, CD127-, CXCR6-, CD25+), and T helper cells (CD3+ CD4+ cells that were neither Tr1 nor Treg). Then, an equal number of each of the suppressor populations was incubated 1:1 with allogenic responder memory CD4+ T cells. The co-cultures were performed in individual wells of a round-bottom 96-well plate, and cells were suspended in X-VIVO^TM^ 15 Serum-free Hematopoietic Cell Media (Lonza Bioscience) supplemented with 10% heat-inactivated human AB serum. Additionally, 1 μg/mL soluble anti-CD3 (OKT3; from Miltenyi) and 3 μg/mL soluble anti-CD28 (CD28.2; from BD) were added as a stimulus to induce proliferation. After 4 days, the cells were stained with the same antibodies used for sorting (as well as anti-CD8a-BV785 [RPA-T8] from BioLegend). 7AAD was used to stain for dying cells immediately prior to analysis by flow cytometry using an Attune NxT Acoustic Focusing Cytometer.

Quantification of proliferation was performed in FlowJo (TreeStar). The percent proliferation was quantified as the number of dimly labelled CD4+ T cells (i.e., CFSE low) divided by the total number of labelled CD4+ T cells (i.e., CFSE+). Percent suppression was calculated as previously described^15^: (“percent proliferation without suppressor” minus “percent proliferation with suppressor”) divided by “percent proliferation without suppressor”.

### Statistics

All analyses were performed in R Version 4.2.2, Stata Version 16, or Python 3. Statistical comparisons of means between groups were performed using two-tailed T tests or the non-parametric Wilcoxon Rank Sum. In instances of repeated measures or when the same biological sample was split between groups, a paired T test was performed. For general comparisons that were agnostic to sample timepoint—for example, comparisons between stimulation conditions (Fig. 5B,D-F, Fig. S5) or between cell population frequencies (Fig. 2, Fig. 5A,C,G-J)—values were averaged across multiple sample timepoints derived from the same individual. When multiple comparisons were made, correction for multiple hypotheses was applied using the Benjamini-Hochberg procedure. These assessments were primarily performed using the Python package pingouin^37^.

Relationships between Tr1 percentages and the odds of symptoms given parasitemia were assessed by logistic regression (Fig. 7A,B). Relationships between Tr1 percentages and the future incidence of malaria was assessed using generalized linear regression models (Fig. 7C,D) or generalized estimating equations with robust standard errors (Table S2). Relationships between Tr1 frequencies and the duration of incident, untreated asymptomatic infections were compared using the non-parametric Wilcoxon Rank Sum Test (Fig. 7E), as well as generalized estimating equations with robust standard errors (Table S2). Repeated measures in participants were accounted for in all models. Multivariable models were adjusted for participant age and log10 transformed parasite density at the time of diagnosis.

## Acknowledgments

Jason Nideffer was supported by a Stanford Interdisciplinary Graduate Fellowship affiliated with Sarafan ChEM-H funded by the philanthropic support of Ruth M. Porat and Anthony Paduano. Florian Bach was supported by a Walter V. and Idun Berry postdoctoral fellowship. Prasanna Jagannathan was supported by the Stanford Maternal and Child Health Research Institute as the Woods Family Faculty Scholar in Pediatric Translational Research. Research was additionally supported by National Institutes of Health grant U01 AI150741 (BG, PJ), National Institutes of Health grant R01 AI177377 (AH, PJ), and Gates Foundation OPP 1113682 (JN, PJ).

## Resource availability

All single-cell genomics data are publicly available from NCBI (BioProject PRJNA1129481). Flow cytometry data is available on request from Prasanna Jagannathan.

