## Supplementary Data for "CXCR6⁺ CD127⁻ Tr1 Cells Balance Immunity and Persistence in Plasmodium falciparum Infection"

**Supplementary Figures**

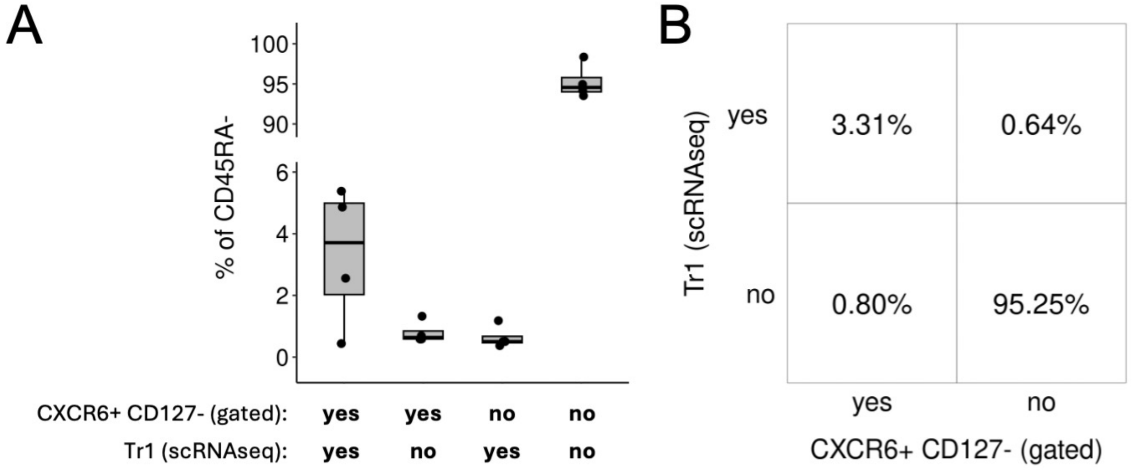


**Fig. S1. Confusion matrix (and underlying data) describing the effectiveness of CXCR6+ CD127- as a gating strategy for identifying Tr1 cells.** (**A**) Box plots displaying the percent of CD45RA- (memory) CD4+ T cells that are Tr1 or non-Tr1 and that fall inside or outside of the CXCR6+ CD127- gate. A break in the y-axis is used to improve data visualization. (**B**) Confusion matrix reporting averages of the percentages in 'A'.

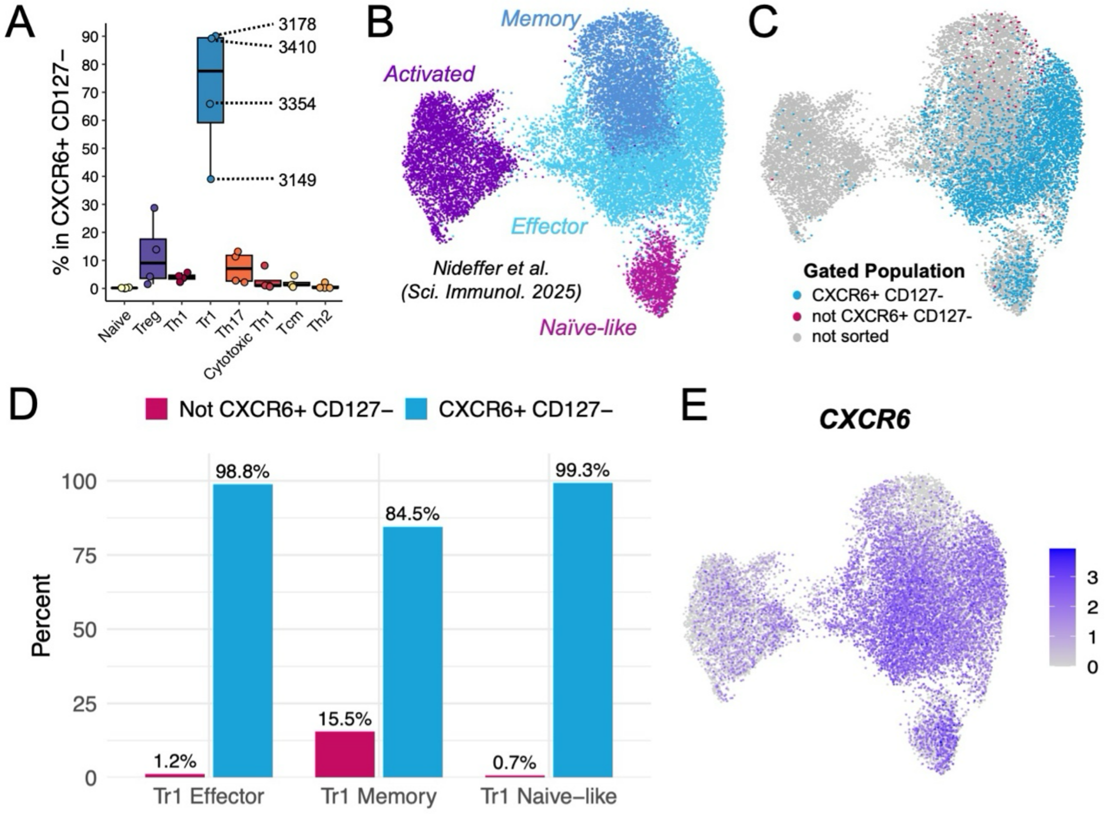


**Fig. S2. Quantification of the heterogeneity amongst CXCR6+ CD127- memory CD4+ T cells.** (**A**) The percentage of CXCR6+ CD127- memory CD4+ T cells that belong to different subsets (defined by scRNAseq). Numbers indicate which data points correspond to each of the four donors. (**B**) UMAP from our prior study depicting the heterogeneity amongst Tr1 cells. (**C**) The same UMAP as in 'B' colored according to whether or not cells were sorted as CXCR6+ CD127-. (**D**) Bar graph quantifying the percentage of a given Tr1 cell subset that was captured (or not captured) by the CXCR6+ CD127- gate. (**E**) The same UMAP as in 'B' colored according to expression of CXCR6.

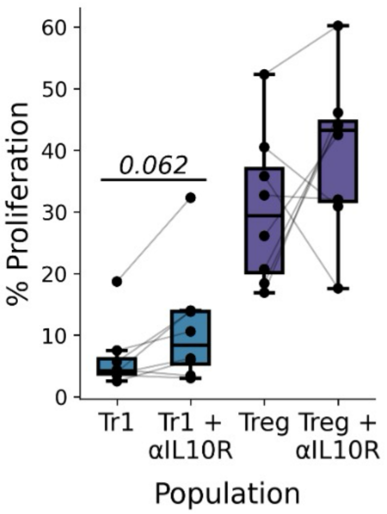


**Fig. S3. Tr1 and conventional Treg proliferation in the presence of IL-10 receptor blockade.**

Cells were incubated for 4 days in the presence of αCD3 and αCD28 as part of a suppression assay. Lines connect different conditions using cells that that were derived from the same donor. Paired T tests were performed comparing conditions with and without IL-10 receptor blockade.

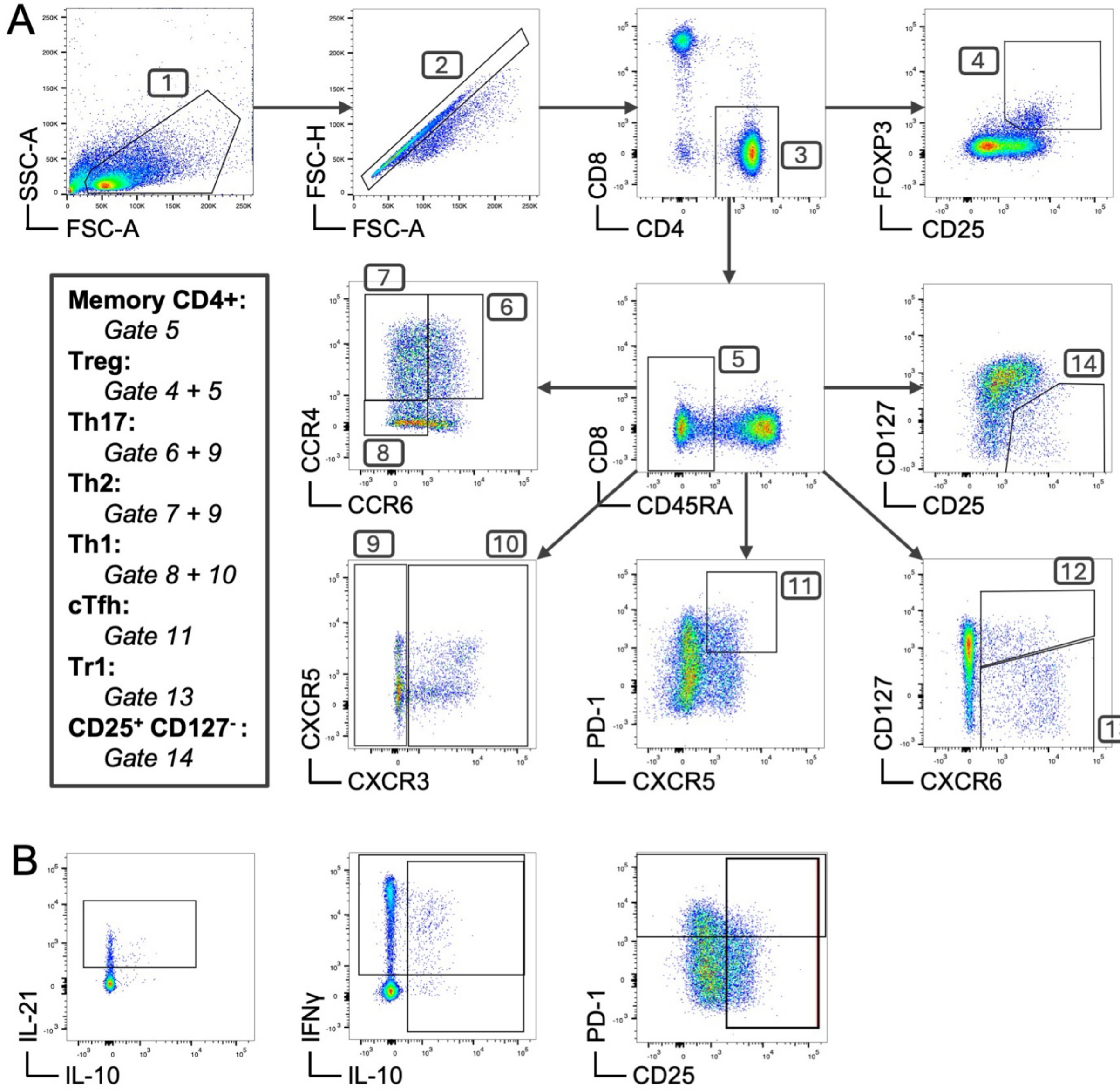


**Fig. S4. Gating of CD4+ T cell subsets and effector molecules.** (**A**) Flow cytometry plots depicting the gating strategy for identifying different cellular populations. (**B**) Flow cytometry plots showing staining of key cytokines and surface-expressed effector molecules.

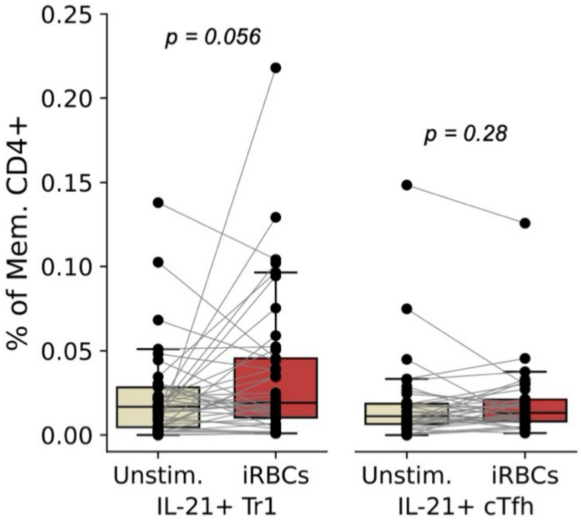


**Fig. S5. IL-21 expression by Tr1 and cTfh cells following stimulation with iRBCs.** The percentage of memory CD4+ T cells that were CXCR6+ CD127- Tr1 (left) or CXCR5+ PD-1+ cTfh and expressed IL-21 in response to iRBC stimulation (compared to unstimulated). Significance was determined via paired T tests.

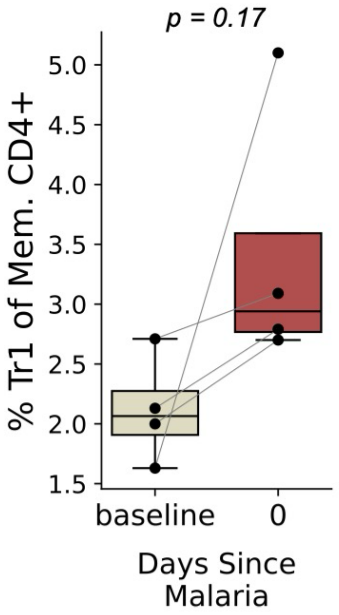


**Fig. S6. Expansion of Tr1 cell frequencies in Ugandan adults diagnosed with malaria.** Tr1 frequencies were measured as a proportion of memory CD4+ T cells at a baseline timepoint prior to infection and at the time of diagnosis. Significance was determined via a paired T test.

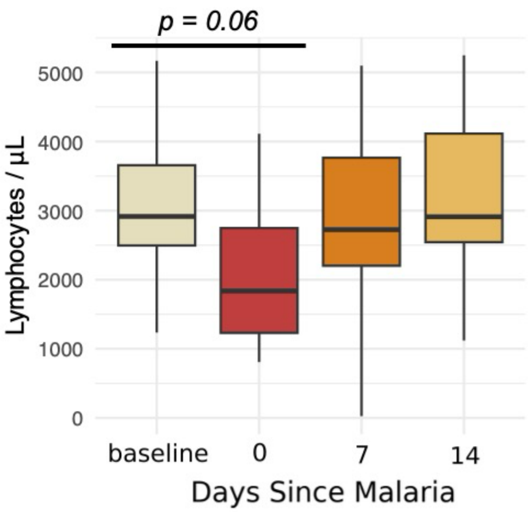


**Fig. S7. Absolute lymphocyte counts in the context of malaria.** Lymphocyte counts were measured before, during, and after symptomatic malaria. Significance was determined using a paired T test comparing mean counts at baseline compared to day 0.

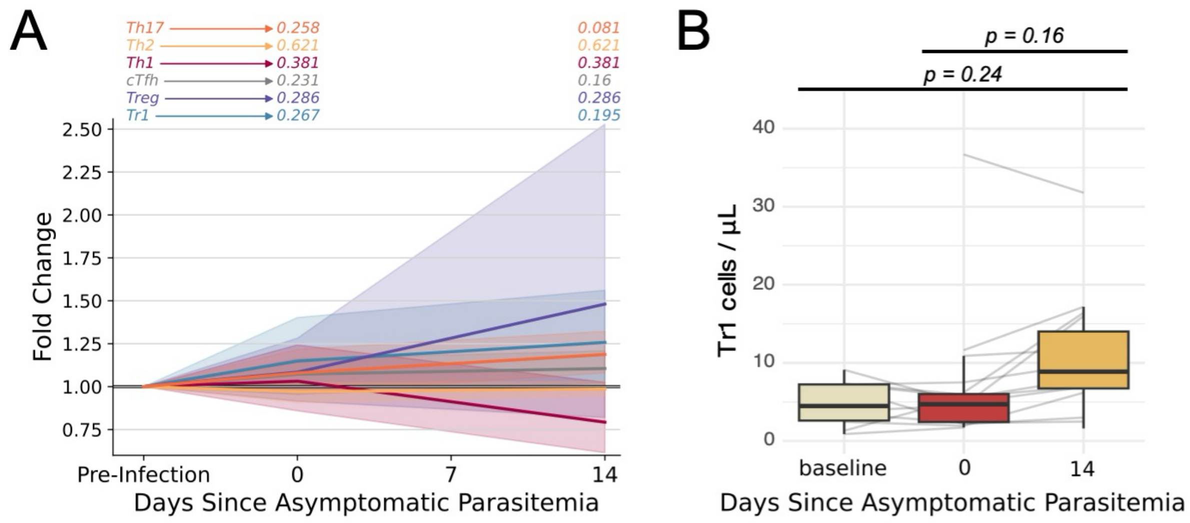


**Fig. S8. Infection dynamics of memory CD4+ T cells populations during asymptomatic parasitemia.** (**A**) Fold change (compared to the pre-infection baseline) in cell frequencies following the detection of asymptomatic parasitemia as determined by flow cytometry. P-values are displayed above each sample timepoint for each population and represent pair-wise comparisons to baseline. (**B**) The absolute counts of Tr1 cells in peripheral blood before, during, and after asymptomatic parasitemia.

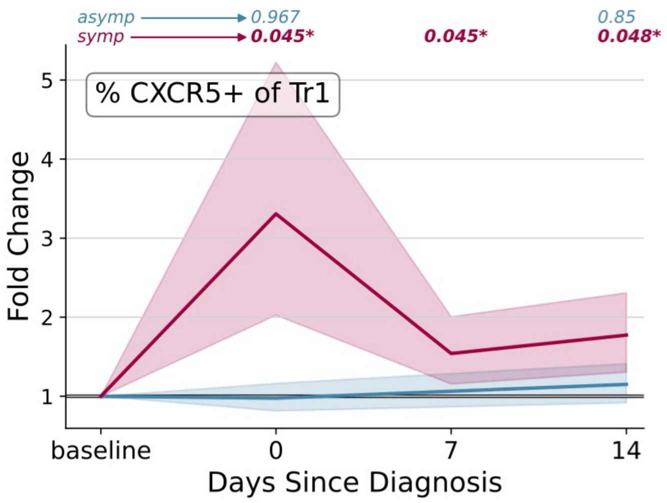


**Fig. S9. Fold change in the percentage of Tr1 cells expressing CXCR5 following symptomatic malaria or asymptomatic parasitemia.** p-values are displayed above each sample timepoint for each infection type and represent pair-wise comparisons to baseline.

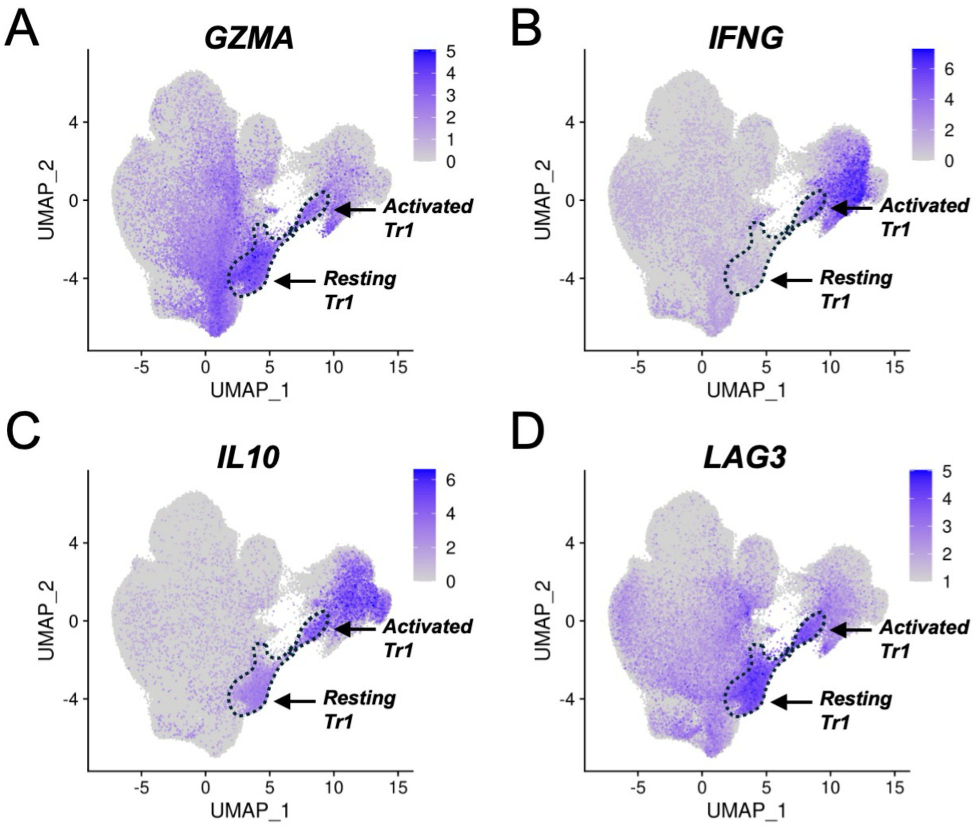


**Fig. S10. Expression of Tr1-associated genes by memory CD4+ T cells.** (**A-D**) UMAPs with the same architecture as Fig. 1A colored according to the expression levels of *GZMA* ('A'), *IFNG* ('B'), *IL10* ('C'), and *LAG3* ('D'). The dashed lines highlight resting and activated Tr1 populations.

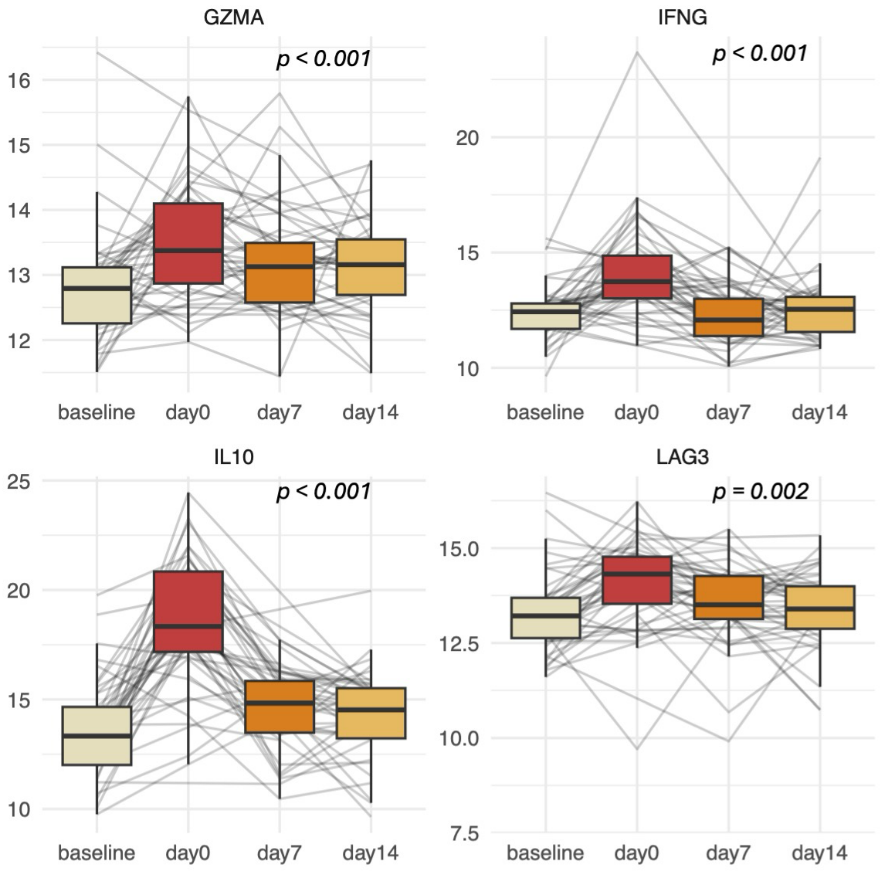


**Fig. S11. Tr1 plasma protein abundance in the context of symptomatic malaria.** The abundance of granzyme-A, IFNγ, IL-10, and LAG-3 in plasma samples (determined by NULISA) collected before, during, and after symptomatic malaria or asymptomatic parasitemia. All p-values shown were adjusted to correct for multiple hypothesis testing. Unless the p-value is explicitly reported, p-value < 0.05 = *; p-value < 0.01 = **; p-value < 0.001 = ***.

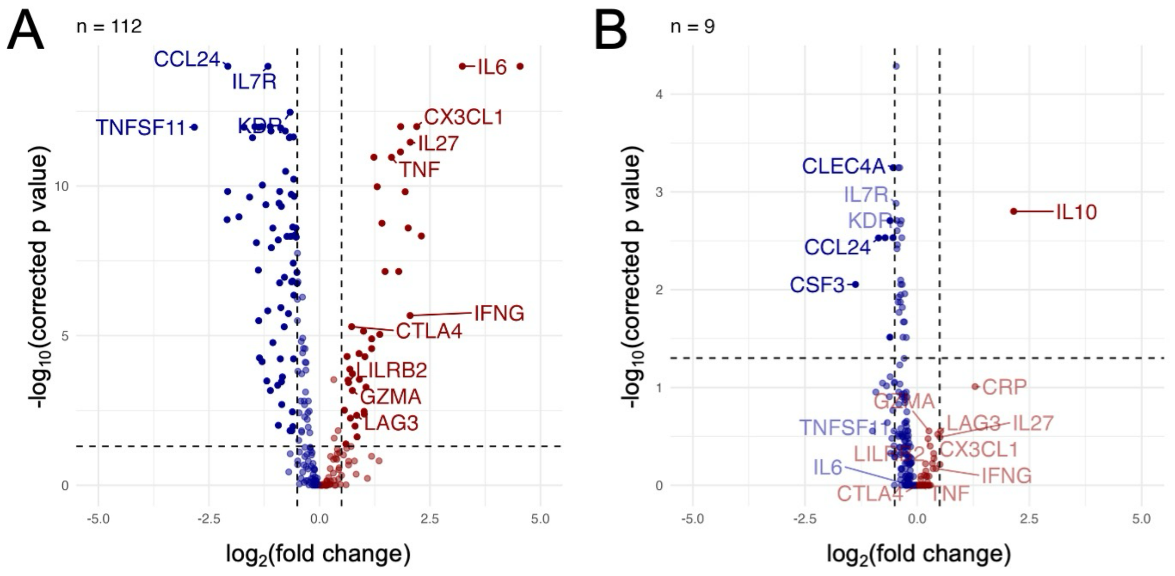


**Fig. S12. Differential abundance of plasma proteins measured by NULISA during symptomatic and asymptomatic infections.** (**A-B**) Volcano plots depicting proteins that are differentially abundant at diagnosis with symptomatic malaria ('A') or asymptomatic parasitemia ('B') versus a pre-infection baseline.

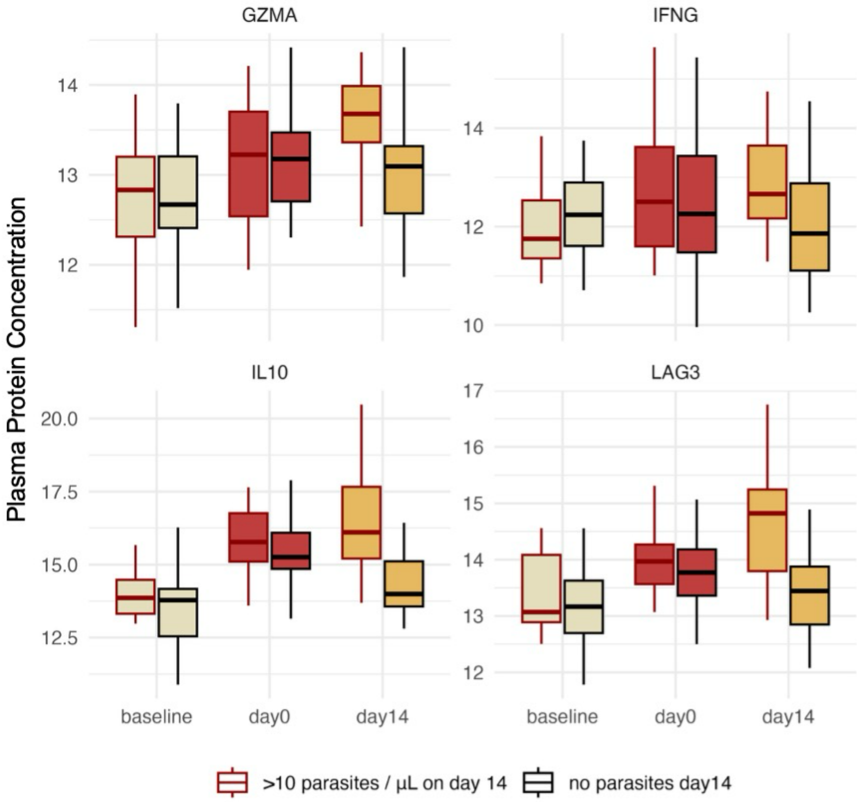


**Fig. S13. The effect of persistent parasitemia on plasma concentrations of Tr1 effector molecules.** The abundance of granzyme-A, IFNγ, IL-10, and LAG-3 in plasma samples (determined by NULISA) collected before during and after asymptomatic parasitemia, stratified by whether the subject tested parasitemia positive or negative at day 14.

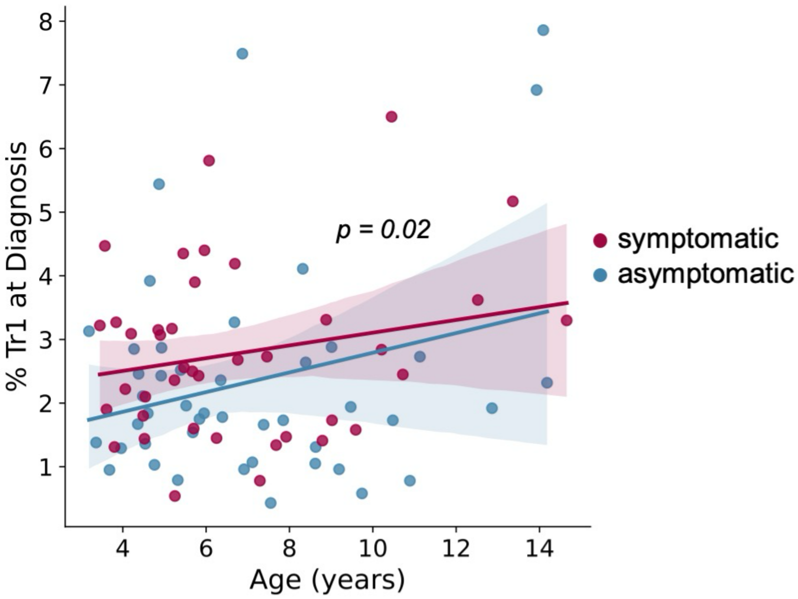


**Fig. S14. Correlation between age and Tr1 frequency at the time of diagnosis.** Separate linear regressions are displayed for symptomatic and asymptomatic infections. Shaded regions represent 95% confidence ranges. The displayed p-value represents the significance of an effect of age on Tr1 frequency in a generalized linear model that includes infection type (symptomatic or asymptomatic) as an independent categorical variable.

**Table S1**. Characteristics of MUSICAL study participants.

| Characteristic | |
|---|---|
| **Age in years at enrollment** | 4.3 (1.8-13.5) |
| **Female sex, n (%)** | 20 (41.7%) |
| **Mean parasite density at time of symptomatic infection, SD** | 18929.2 (19955.39) |
| **Mean parasite density at time of asymptomatic infection, SD** | 3453.3 (6038.5) |
| **Sickle cell status*** | |
| **HbAA, n/N(%)** | 36/46 (75.0%) |
| **HbAS, n/N(%)** | 10/46 (20.8%) |
| **Incidence of malaria per person year over period of cohort (2020-2023)** | 1.96 |

*2 individuals with missing Hb genotype data

**Table S2**. Univariate and multivariate analyses of the relationship between Tr1 frequencies and the future incidence of malaria or the duration of incident asymptomatic infections.

| | Future incidence of symptomatic malaria over 2 years | | | | Duration of incident asymptomatic infection in days[1] | | | |
|---|---|---|---|---|---|---|---|---|
| | Univariate | | Multivariate[2] | | Univariate | | Multivariate[2] | |
| | IRR (95% CI) | P | IRR (95% CI) | P | Coef (95% CI) | P | Coef (95% CI) | P |
| **Age in years** | 0.89 (0.80-0.98) | 0.02 | 0.87 (0.78-0.97) | 0.01 | 0.38 (-3.4-4.2) | 0.84 | 2.2 (-2.5-6.9) | 0.36 |
| **Log10 parasite density at diagnosis** | 1.06 (0.82-1.37) | 0.67 | 1.02 (0.78-1.32) | 0.89 | 4.15 (-6.9-15.2) | 0.46 | 7.36 (-1.9 – 16.6) | 0.12 |
| **%CXCR6+CD127- Tr1 prior to infection** | | | | | | | | |
| **Group 1, n=22 (0.43%-1.5%)** | Ref | Ref | Ref | Ref | Ref | Ref | Ref | Ref |
| **Group 2, n=40 (1.5%-3.08%)** | 1.43 (0.83-2.48) | 0.20 | 1.39 (0.76-2.55) | 0.29 | 23.7 (-3.2-50.6) | 0.085 | 27.3 (-3.5-58.2) | 0.08 |
| **Group 3, n=23 (3.1% - 7.86%)** | 0.77 (0.37-1.59) | 0.48 | 0.76 (0.37-1.59) | 0.47 | 33.0 (-6.9-3.0) | 0.10 | 42.5 (3.6-81.4) | 0.032 |
| **%CXCR6+CD127- Tr1 at time of diagnosis** | | | | | | | | |
| **Group 1, n=22 (0.43%-1.5%)** | Ref | Ref | Ref | Ref | Ref | Ref | Ref | Ref |
| **Group 2, n=40 (1.5%-3.08%)** | 0.74 (0.33-1.62) | 0.45 | 0.74 (0.35-1.57) | 0.44 | 15.5 (-14-45.1) | 0.31 | 11.23 (-21.5-44.0) | 0.50 |
| **Group 3, n=23 (3.1% - 7.86%)** | 0.38 (0.18-0.80) | 0.011 | 0.38 (0.18-0.80) | 0.011 | 51.8 (17.4-86.3) | 0.003 | 52.67 (18.9-86.5) | 0.002 |

IQR:
[1]Incidence infections defined by AMA1 amplicon sequencing.
[2]Multivariate models adjusting for age in years and $log_{10}$ parasite densities (detected by quantitative PCR)
